# MMP14 cleaves PTH1R in the chondrocyte derived osteoblast lineage, curbing signaling intensity for proper bone anabolism

**DOI:** 10.1101/2022.08.17.504252

**Authors:** Tsz Long Chu, Peikai Chen, Anna Xiaodan Yu, Mingpeng Kong, Zhijia Tan, Kwok Yeung Tsang, Zhongjun Zhou, Kathryn S.E. Cheah

## Abstract

Bone homeostasis is regulated by hormones such as parathyroid hormone (PTH). While PTH can stimulate osteo-progenitor expansion and bone synthesis, how PTH-signaling intensity in progenitors is controlled is unclear. Endochondral bone osteoblasts arise from perichondrium-derived osteoprogenitors and hypertrophic chondrocytes (HC). We found, via single-cell transcriptomics, HC descendent cells activate membrane-type 1 metalloproteinase 14 (MMP14) and the PTH pathway as they transition to osteoblasts in neonatal and adult mice. Unlike *Mmp14* global knockouts, HC lineage-specific *Mmp14* null mutants (*Mmp14^ΔHC^*) produce more bone. Mechanistically, MMP14 cleaves the extracellular domain of PTH1R, dampening PTH signaling and PTH signaling is enhanced in *Mmp14^ΔHC^* mutants. We found HC-derived osteoblasts contribute ∼50% of osteogenesis promoted by treatment with PTH 1-34 and this response was amplified in *Mmp14^ΔHC^*. MMP14 control of PTH signaling likely applies also to both HC- and non- HC-derived osteoblasts because their transcriptomes are highly similar. Our study identifies a novel paradigm of MMP14 activity-mediated titration of PTH signaling in the osteoblast lineage, contributing new insights into bone metabolism with therapeutic significance for bone-wasting diseases.

## Introduction

Developing and maintaining appropriate proportions of cells of the osteoblast lineage and other cell types (e.g., osteoclasts, endothelial cells, adipocytes, stem cells) in the bone marrow are crucial for healthy bones. These cell types together maintain bone mass via concerted activities of bone building (anabolic) by osteoblasts versus bone resorption (catabolic) by osteoclasts, with input from factors produced by osteocytes to orchestrate the remodeling process (1). Through this continuous remodeling, and balanced anabolic and catabolic activity, a homeostatic condition is achieved (2). This homeostasis is regulated by growth factors and cytokines produced within bone and systemic factors such as parathyroid hormone (PTH) and estrogens (3, 4). An imbalance in proportions of these cells in bone impairs bone resorption and formation, which can result in diseases of bone mass such as osteopenia, osteoporosis, and osteopetrosis (5, 6).

Long bones form by endochondral ossification, a process in which chondrocytes differentiate, proliferate, mature and become hypertrophic, forming a cartilaginous growth plate that coordinates longitudinal bone growth and is recapitulated in fracture repair (7, 8). Hypertrophic cartilage is remodeled by matrix metalloproteinases (MMPs) (reviewed in(9). The chondrocyte differentiation cascade is coordinated by many transcription factors and signaling pathways, including reciprocal signaling via the parathyroid hormone-related protein (PTHRP)-Indian hedgehog (IHH) feedback loop (10, 11). Resting chondrocytes secrete PTHRP, that binds to parathyroid hormone 1 receptor (PTH1R) expressed by pre-hypertrophic chondrocytes, acting to delay their differentiation to hypertrophic chondrocytes (HCs) (12, 13). HCs specifically produce collagen type X (*Col10a1*), down-regulate PTH1R and undergo distinct phases of cell enlargement for bone elongation (14, 15). Recent work has shown that osteoblasts in trabecular bone are derived from HCs in the growth plate and from osteoprogenitors in the perichondrium which accompany invading blood vessels (15–17). HCs have been shown to contribute to the full spectrum of cells in the osteoblast lineage in endochondral bone, trabeculae, endosteal/endocortical bone, including osteocytes, and, to a minor degree, bone marrow stromal cells and adipocytes (15, 17–20). HC transformation to osteoblasts also occurs in bone healing (7, 15, 17). The functional importance of the HC lineage has been demonstrated in mice, which have reduced bone mass as a consequence of ablating β-catenin and *Irx3/5* genes specifically in HCs (19, 21). However, it is not known whether the HC-derived osteogenic lineage is molecularly distinct from the non-HC lineage (contributed by the perichondrium, periosteum, bone marrow mesenchymal stem cells), which specific MMPs act at the chondro-osseous junction to facilitate the transition of HCs to osteoblasts and what physiological contribution(s) they make to maintaining bone homeostasis and anabolism in response to extrinsic signals.

Here, using single-cell transcriptomics, we characterized the different osteoblast populations in endochondral bone and found that their transcriptomes are broadly similar to that of non-HC derived osteoblasts. We found HC-derived osteoblasts activate membrane-type 1 metalloproteinase 14 (MMP14) and the PTH/PTH1R pathway upon transition into sub-chondral bone. Using mouse mutants and biochemical approaches, we identified the cleavage of PTH1R by MMP14 in HC-derived osteoblasts as a novel mechanism that curbs the intensity of PTH signaling. Like their non–HC-derived counterparts, HC-derived osteoblasts respond to exogenous PTH treatment, and contribute significantly to the bone anabolic response. MMP14 modulation of PTH signaling intensity controls the differentiation and survival of osteoblasts and thereby the anabolic response to PTH in building bone mass.

## Results

### HC-derivatives activate MMP14 as they transition to become osteoblasts

To reveal key transcriptional changes in HC lineage cells as they translocate across the chondro osseus junction to contribute to forming trabecular bone, we exploited *Col10a1-Cre* (*C10Cre*); *Rosa26-tdTomato* (*RtdT*) (abbreviated *C10Cre;RtdT*) mice in which the HC-derived osteoblast lineage is marked by expression of tdTomato because of the specific activity of Cre recombinase on the *RtdT* Cre reporter in HCs (15, 19, 22). We sequenced the RNA of single-cells (scRNAseq) isolated from the tibia of *C10Cre;Irx3^+/ΔHC^Irx5^+/-^;R26^td/+^* postnatal day 6 (P6) (phenotypically normal, (19)) and *C10Cre;RtdT* 8-week (P56) mice (Fig. 1A; Methods). For P6 tibia single cells, 12 clusters were identified by dimension reduction analyses (Fig. 1B and Methods). Of these, 8 clusters expressed genes characteristic of chondro-osteogenic cells, which we annotated by reference to the literature (23–25) as resting (e.g. *Pthlh)*, proliferating (e.g. *Top2a, Mki67)*, and maturing chondrocytes (e.g. *Col9a2, Matn1*), pre-hypertrophic chondrocytes (pre-HCs)(e.g. *Ihh, Fgfr3, Pth1r*), HCs (*Col10a1*), immature (*Mmp14, Pdgfrb)* and mature osteoblasts (*Ifitm5, Bglap*), and osteocytes, (corresponding to terminally differentiated osteoblasts, *Sost1, Dmp1, Phex*), respectively (Fig. 1B, Fig. S1A, and Methods).

**Figure 1.**
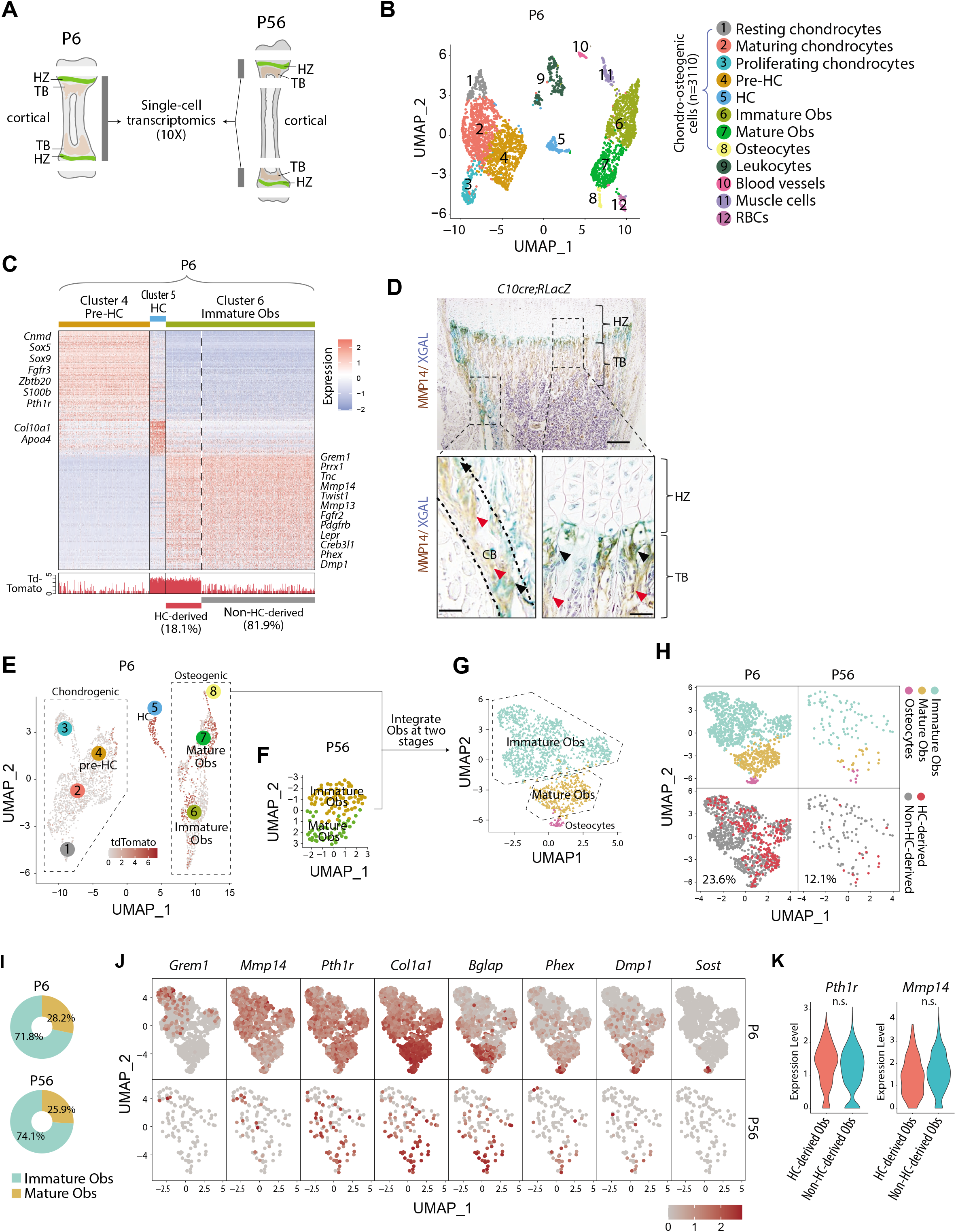
Single-cell transcriptomics analyses of endochondral bone sub-populations at P6 and P56. (A), Schematic diagram showing the isolation of osteochondrogenic cells from mouse tibia for single-cell transcriptomics (scRNAseq). (B), Scatter-plot showing the 3420 single cells, 3110 of which belong to the chondro-osteogenic clusters, as identified in the P6 sample by dimension reduction method UMAP. HC: hypertrophic chondrocytes. Obs: osteoblasts. RBC: red blood cells. (C), Upper panel: heatmap showing the expression patterns of P6 signature genes in the preHC, HC and immature Ob clusters, when comparing in a one-vs-all manner among the three. Representative signature genes of interest are listed by the side. Lower panel: signal track indicating the level of tdTomato expression inferred from the WPRE signal. (D), Representative immunostaining showing the spatial distribution of MMP14 and X-gal stained (blue) in hematoxylin stained sections from C10Cre;RLacZ tibae. HZ, hypertrophic zone; TB, trabecular region; CB, cortical region. Black arrows in enlarged figures indicate LacZ+;MMP14+ cells. Red arrows indicate Mmp14+;LacZ-cells. Dotted line draws chondro-osseous junction. Scale bar 200μm,50μm(magnified picture).(E-H), Scatter-plot showing the integration of the mature and immature osteoblasts of the P6 (E) and the P56 (F) mice. (H-J), (I), Piecharts showing the percentages of mature osteoblasts in the both samples. (J), A panel of scatter-plots showing the expression pattern of a list of selected markers, representing the different osteoblast differentiation states. See also Figure S1, Extended Table S1&2.

We identified osteoblasts as HC-derived if they transcribed tdTomato, using the alignment to the tdTomato sequence and a surrogate sequence for it (WPRE (26, 27), Methods, Fig. S1, A, E-G). We annotated the different cell populations based on the hundreds of differentially expressed signature genes (Extended-Data Table 1) associated with each cell population and by reference to the literature (Fig. S1). Immature osteoblasts represent an important early stage in the HC to osteoblast transition and 18.1% of them were found to express tdTomato and thus were annotated as HC-derived (Fig. 1C and Fig. S1H). We used well established signature genes (23–25) such as *Pth1r, Col10a1* and *Grem1* to annotate the different populations as Pre-HCs, HCs and immature osteoblasts respectively (Fig. 1C, see also Methods and Extended Data Table 2). Gene-ontology (GO) enrichment analyses suggests a series of developmental events consistent with onset of endochondral ossification (Fig. S1B-D and Extended-Data Table 2) (16) and confirms the lineage continuum of HCs and osteoblasts.

Given the need for HCs to be released from the cartilage matrix as they move across the chondro-osseous junction, MMP genes are candidates, especially those expressed at that region and in the adjacent sub-chondral bone. Amongst the *Mmps, Mmp13* was expressed in both the HC and immature Ob populations ((15, 23) Fig. 1C, Extended-Data Table 1), consistent with the literature (28). By contrast, *Mmp14* (encoding a membrane-anchored MMP (29) was not expressed in HCs but was enriched in immature osteoblasts (Fig. 1C, Fig.S1A, Extended-Data Table 1). Consistent with the scRNAseq results, immunostaining for MMP14 detected no protein in HCs and strong expression in LacZ-labeled HC-descendant cells, especially those located at the chondro-osseous junction (Fig. 1D) (identified by co-staining with LacZ in *C10Cre*; *Rosa26-LacZ* mice (15)).

We next investigated whether the molecular characteristics of the osteogenic signatures in HC-derived osteoblasts persisted by maturity at P56 (Fig. 1, E-J and Fig. S1, I, J). We integrated the osteogenic cell clusters in the P6 (Fig. 1E) and P56 (Fig. 1F) samples. Although fewer osteogenic cells were isolated in the P56 sample compared to P6, the overall relative proportions of immature osteoblasts were similar between P6 and P56 (28.2% and 25.9%, respectively) (Fig. 1I). We found 94.7% of the immature osteoblasts (expressing *Grem1, Mmp14)* and 63.4% of the mature osteoblasts (expressing *Col1a1, Bglap)* and osteocytes (expressing *Phex, Dmp1, Sost*) at P56 were correctly mapped to the relevant clusters at P6 (Fig. 1, G-J; Extended-Data Table 1). Using the same cutoff for tdTomato expression, 23.6% of the P6 and 12.1% of the P56 osteogenic cells (combining immature and mature Obs) were HC-derived (Fig. 1H). Overall, the HC-derived osteogenic populations in young post-natal and mature mice are highly comparable in molecular signatures, cell-type and lineage composition, consistent with a hypertrophic origin of osteogenic cells (Fig. 1J). The variations in relative percentages may also be related to the limits in sensitivity in detecting tdTomato mRNAs and differences in the ease of releasing deeply embedded cells within the bone matrix versus more superficially located ones. However these frequencies are in agreement with the broad ranges (18-60%) reported previously for HC-derived osteoblasts from a range of developmental and postnatal ages (15, 17, 30).

We also assessed how similar HC-derived and non-HC-derived cells are. While HC-derived and non HC-derived osteogenic cells generally overlapped (Fig. 1E, H and Fig. S1, I), the HC-derived cells tended to localize in certain regions in the integrated UMAP plot (Fig. 1H and Fig. S1I), suggesting there may be subtle transcriptomic differences between the two lineage origins. Among the 560 genes uniquely up-regulated in immature osteoblasts, only 22 are differentially expressed between HC- and non-HC-derived cells (Fig. S1J). This suggests a high degree of similarity between these two sources of immature osteoblasts. Interestingly, both sources expressed progenitor cell (*Lepr* and *Grem1* (31) and mesenchymal cell (*Prrx1*, *Twist1, Pdgfrb*) markers (Fig. 1C), which is consistent with a recent report (30) suggesting these cells pass through progenitor and mesenchymal states.

Comparing the immature and mature HC- and non-HC-derived osteoblasts combined, we detected 48 differentially expressed genes, of which 22 were expressed higher in the HC-derived cells, including some mature osteoblast markers, such as *Ifitm5*, *Col22a1*, and *Smpd3*; and 26 genes were expressed higher in the non-HC-derived cells, such as *Postn*, *Igfbp5*, *Col3al* (Fig. S1J). This small set of differentially expressed genes may reflect heterogeneity and differences in maturities between osteoblasts of different origins, the significance of which will be investigated in future studies. Overall, there was a broad similarity in transcriptomic characteristics for the HC-derived and non-HC-derived osteoblasts.

### Global MMP14 is required for proper translocation of HC-derivatives to trabecular bone

*Mmp13* and *Mmp14* are candidate facilitators for the translocation of HCs to the subchondral space. *Mmp13* is expressed in late HCs (15, 23) as well as osteoblasts. Chondrocyte-specific *Mmp13* knockout mice display abnormal growth plates with disrupted terminal hypertrophy and increased trabecular bone (28), which could reflect an impact on the HC-Ob lineage continuum. By contrast MMP14 is expressed specifically at the chondro-osseous junction in immature HC-derived osteoblasts but not in HCs themselves (Fig. 1, C, D). Amongst MMPs, *Mmp14* null mutants display the most severe skeletal phenotypes, including loss of trabecular bone, over-activity of osteoclasts, impaired angiogenesis and reduced calvarial ossification (29, 32–34), consistent with expression of *Mmp14* by many cell types, such as skeletal progenitors, osteoblasts, osteoclasts, endothelial cells and bone marrow stromal cells (Fig. S2A) (34–38). The impaired bone formation in *Mmp14*-deficient mice and the expression of MMP14 in HC-derived osteoblasts raised the possibility that MMP14 regulates their differentiation. The impact of chondrocyte-specific knockout of *Mmp14* on trabecular bone formation is not known. Given MMP14’s pivotal role in trabecular bone formation, we focused on *Mmp14* and asked whether dysfunction in lineage progression of HC-derived cells was responsible for the severe bone deficit in *Mmp14* knockout mice. We used the *C10Cre;Rosa26-YFP* reporter to lineage trace HC-derivatives in global *Mmp14* knockout mice, and found an increased accumulation of HC-descendants at the chondro-osseous junction compared to WT controls (Fig. S2B and C). There was also a small increase in proliferating HC-derivatives (marked by 5-Ethynyl-2’-deoxyuridine, Edu) in the region immediately below the chondro-osseous junction (Fig. S2C). However there were fewer proliferating HC-derivatives in further distal regions below the chondro-osseous junction, consistent with their accumulation there and impaired translocation into the trabecular bone (Fig. S2C).

### Intrinsic MMP14 activity in HC-derivatives is not essential for their translocation to trabecular bone

To investigate whether the stagnation of HC-descendants under the chondro-osseous junction reflected impaired translocation to trabecular bone in *Mmp14* global knockouts and was intrinsic to HC-derived cells themselves or involved extrinsic influences from other cell types, we genetically inactivated *Mmp14* in HC-descendants by generating HC-specific conditional *Mmp14^Flox^ (Mmp14^F/F^)* mutants using *C10Cre* (abbreviated as *Mmp14^ΔHC^*) (Fig. 2A). We found removing *Mmp14* in HC-descendants did not recapitulate the accumulation of HC-descendants at the chondro-osseous junction observed in *Mmp14^-/-^* mice (Fig. S2C). We used tamoxifen mediated pulse-labelling and chasing of *Mmp14* deficient HC-descendants using *C10Cre-ERT;Mmp14^F/F^;RtdT* to assay the dynamics of their translocation into the subchondral space. We found the localization and distribution of HC-descendants were unaffected in *Mmp14^ΔHC^* mice (Fig. 2B). These results suggest the aberrant stagnation of HC-derivatives at the chondro-osseous junction was not intrinsic to a defect in these cells themselves but might be a consequence of MMP14 deficiency elsewhere, for example in the invading vascular cells and/or the accompanying osteoprogenitors from the perichondrium that co-migrate with blood vessels to populate the primary spongiosa (16, 39). Vascular invasion is required for bone formation (39, 40). We therefore examined the vascular capillaries in *Mmp14^-/-^* mice and found that both vascular density and endothelial cell count (measured using the marker ENDOMUCIN) were decreased compared with control mice (Fig. S2D). In vitro and in vivo studies have shown *Mmp14* is important for angiogenesis, neovessel formation and migration (29, 41). The abnormal vascularization in *Mmp14* null mice may therefore be a major contributor to the compromised translocation of HC-derived cells to the trabecular bone.

**Figure 2.**
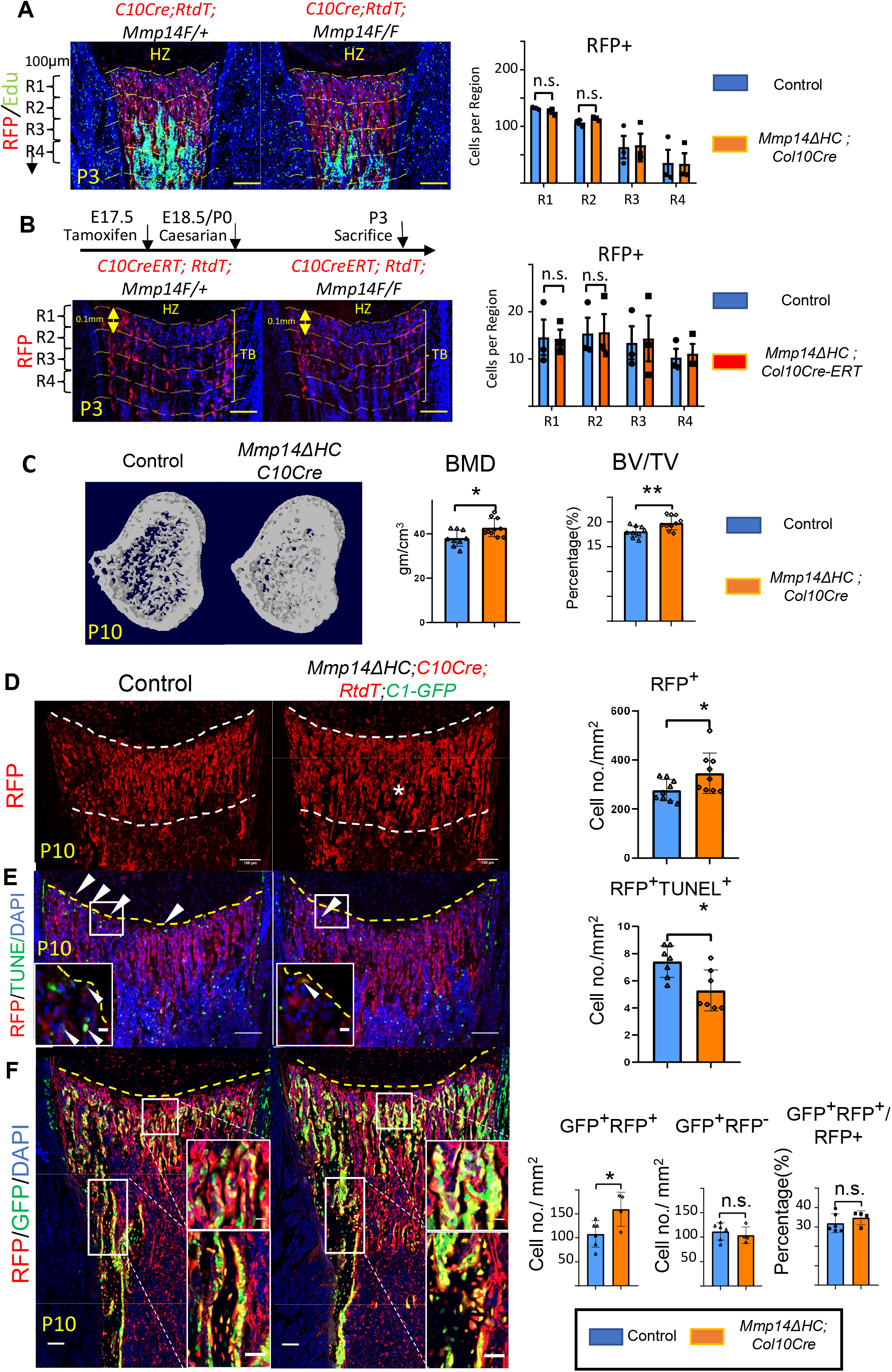
Increased osteogenesis in Mmp14ΔHC is not because of abnormal transition of HCs. (**A**), Immunofluorescence staining of RFP and Edu-labeled cells counterstained with DAPI in Mmp14ΔHC (orange bar) and control mice tibia at P3. For quantification, the trabecular bone is divided into four zones, each 0.1 mm in thickness. The number of RFP+(red) cells in each region were counted. (**B**), Schematic of experimental design. Tamoxifen was administered to C10CreERT;RtdT;Mmp14F/+ and C10CreERT;RtdT;Mmp14F/F mice at E17.5 and harvested at P3. Staining and quantification as (**A**). (**C**), Transverse image of Micro-CT analysis in Mmp14ΔHC mutants and their littermate controls at P10. Statistical analysis of bone mineral density(BMD) and bone volume over tissue volume ratio(BV/TV) using Micro-CT in Mmp14ΔHC and their littermate control(blue bar) (n=9). The data points were pooled across multiple bone samples. (**D**), Representative immunofluorescence staining and quantification of tdTomato labeled HC-descendants in trabecular bone (asterisk) of Mmp14ΔHC and littermate controls at P10. White dotted line highlights the region below the chondro-osseous junction. Scale bars, 100 μm. The data represent means ± SEM. The data points were pooled across multiple bone samples (n = 7). (**E**), *In situ* terminal deoxynucleotidytransferase deoxyuridine triphosphate nick end labeling(TUNEL) assay labeling apoptotic cells and quantification in Mmp14ΔHC and control mice at chondro-osseous junction at P10. (n=7). (**F**), Representative immunofluorescence staining and quantification of RFP(red), GFP(green) counterstained with DAPI(blue) marking tdTomato labeled HC-descendants, GFP labeled osteoblasts and nucleus in Mmp14ΔHC;RtdT;Col1-2.3-GFP(C1-GFP) and control littermates at P10. RFP+GFP+, RFP-GFP+ and ratio of GFP+RFP+/RFP+ cells in the trabecular bone comparing Mmp14ΔHC and littermates. The data points were pooled measurements across multiple bone sections per animal. n = 4 for Mmp14ΔHC, n=6 for control. Statistics: The data represent means ± SEM. Fig. b, e, g, h: p values were calculated using two-tailed unpaired t-tests, * p<0.05, ** p<0.01, *** p<0.001.

### Increased osteogenesis and number of HC-descendants in *Mmp14 ^ΔHC^* mutants

To delineate the intrinsic role of *Mmp14* in the HC-lineage and on trabecular bone formation, we characterized the bone phenotype of *Mmp14^ΔHC^* mice. Interestingly, MicroCT analysis revealed a 16% increase in BMD and a 10% increase in trabecular BV/TV in *Mmp14^ΔHC^* mice (Fig. 2C). The increased bone mass in *Mmp14^ΔHC^* mutants, in contrast to *Mmp14^-/-^*, led us to ask whether the increased BMD was a consequence of increased osteogenesis. To elucidate whether increased trabecular bone density and volume in *Mmp14^ΔHC^* mice is directly attributable to osteogenic HC-descendants, we addressed the fate of HC-descendants at postnatal stages (Fig. 2D-F). Immunofluorescence staining of MMP14 and RFP shows the number of HC-descendants was increased in *Mmp14^ΔHC^* mice at P10 (Fig. 2D). This observation could be due to a decrease of TUNEL+ cells at the chondro-osseous junction (Fig. 2E). By using *Col1a1-2.3-GFP* (*C1-GFP*) to co-label osteoblasts, we observed an increased number of HC-derived osteoblasts in *Mmp14^ΔHC^* mutants, compared to similar GFP+RFP-cell counts between mutants and controls (Fig. 2F). The data collectively suggests targeting *Mmp14* in HCs promoted osteogenesis. Consistent with the above findings, the expression domains of osteoblast markers *Col1a1* and *Mmp13* were expanded in *Mmp14^ΔHC^* mice (Fig. S3A). In contrast to reduced apoptotic cells, the number of Edu+ labeled proliferating cells was comparable between *Mmp14^ΔHC^* mutants and control mice (Fig. S3B).

To test if the number of osteoclasts was affected in *Mmp14^ΔHC^* mutants, we performed Tartrate-resistant acid phosphatase (TRAP) staining and quantitated the number of osteoclasts in trabecular bone (Fig. S3C). In line with reports showing MMP14 deficiency does not affect osteoclastogenesis (42, 43), the number of osteoclasts was comparable between *Mmp14^ΔHC^* mutants and controls at P10, suggesting the increased number of HC-derived osteogenic cells, not reduced resorption, was a major contributor for increased bone mass at this stage. Overall these results are in line with previous reports showing a combined inactivation of MMPs is required for osteoclast dysfunction (43).

A small proportion (less than 2%) of WT HC-descendants have been shown to become bone marrow adipocytes (19, 20). We found the total frequency of HC-derived bone marrow adipocytes in *Mmp14^ΔHC^* mice was unchanged compared with control, suggesting MMP14 does not influence the adipogenic fate choice of HC-derived cells. However non-HC derived adipocytes in *Mmp14^ΔHC^* were substantially fewer than in controls, suggesting a non-cell-autonomous impact of MMP14 deficiency on signals from HC-derived cells on marrow adipogenesis in *Mmp14^ΔHC^* mice (Fig. S3D). Such non cell-autonomous effects could be mediated by the effect of MMP14 deficiency on the collagenous microenvironment that coordinates adipogenesis (44, 45).

### PTH1R is a substrate of MMP14

Next we sought to determine the underlying reason for the increased bone in *Mmp14^ΔHC^* mice. Reduced calvarial osteogenesis and osteoclast overactivity in *Mmp14* mutant mice were found attributable to cleavage of ADAM9, RANKL and extracellular matrix proteins by MMP14, regulating FGF, RANK and YAP/TAZ signaling (33, 34, 42). However, these findings are insufficient to account for the increased trabecular bone observed in *Mmp14^ΔHC^* mice, suggesting a yet undiscovered mechanism could be the underlying cause of the phenotype. Interestingly, MMP14 was shown to be a downstream target of PTH signaling in osteocytes (46). In a previous report, because cleavage of PTH1R can be inhibited by TIMP2, but not TIMP1 it was proposed that MMP-dependent cleavage of PTH1R causes reduced stability and degradation of PTH1R, raising the possibility that cleavage by a MMP might inhibit PTH signaling (47). However the identity of the responsible MMP was unknown. This collective evidence, including the co-expression of *Mmp14* and *Pth1r* from scRNA data (Fig. 3A), strongly suggested a direct molecular link between MMP14 and PTH/PTH1R signaling pathway could be possible. Therefore, we hypothesized MMP14 can proteolytically process and perhaps inhibit PTH/PTH1R signaling. Knocking out MMP14 in HC-derived osteogenic progeny could remove its inhibitory effect on PTH pathway.

**Figure 3.**
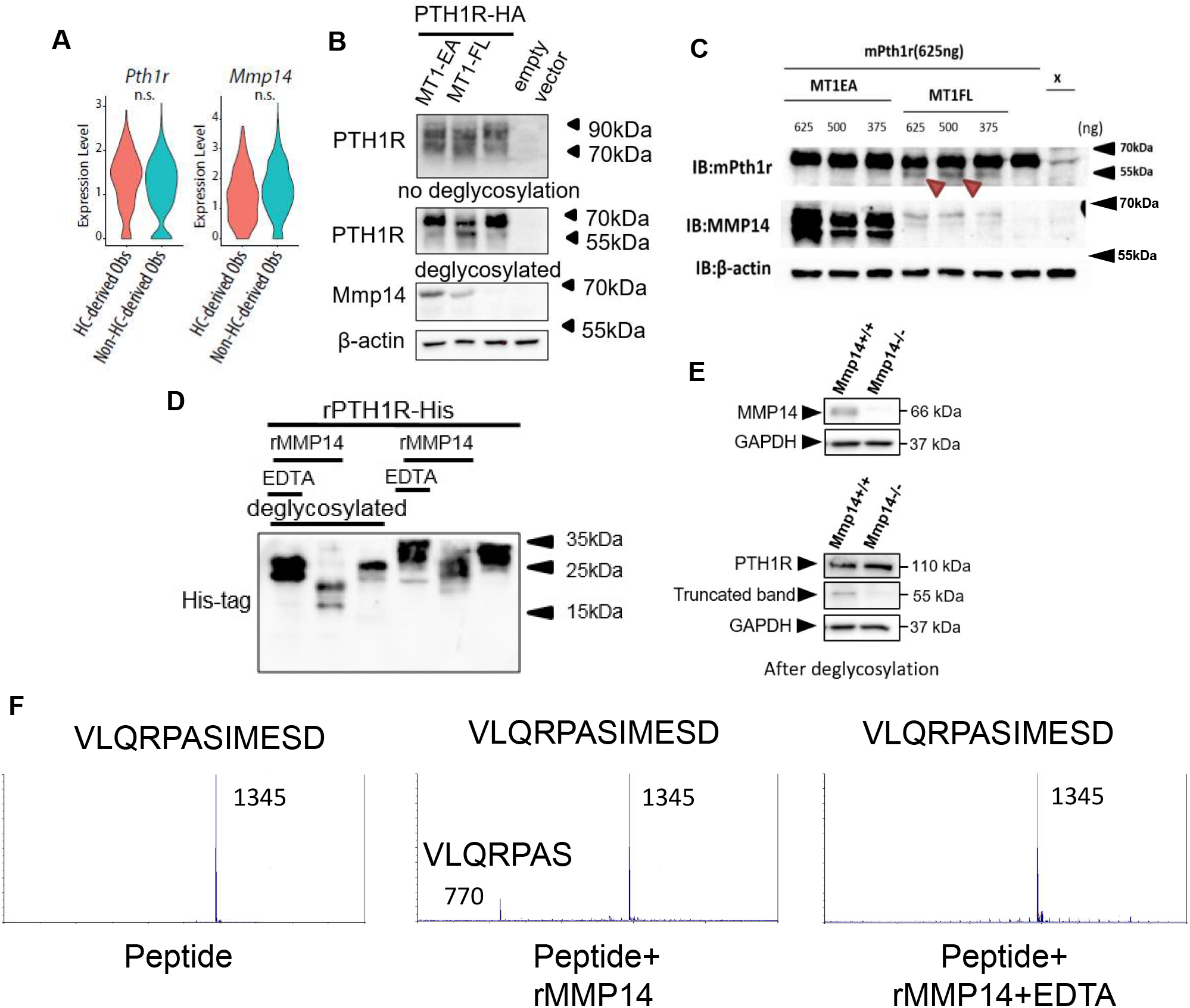
PTH1R is a substrate of MMP14. (**A**), Violin plots showing the expression levels of Pth1r and Mmp14 in the HC-derived and non-HC-derived osteoblasts. n.s.: not significant. **(B)**, HEK293T cells expressing HA-tagged human PTH1R were transfected with either WT(MT1-FL) or E/A catalytic inactive mutant MMP14 (MT1 EA). β-actin is used as a loading control. Image representative of three independent experiments. (**C**), HEK293T cells expressing mouse Pth1r(without HA-tag) were transfected with either WT(MT1-FL) or E/A catalytic inactive mutant MMP14 (MT1-EA). Mouse Pth1r was deglycosylated, detected with anti-PTH1R antibody and β-actin is used as a internal loading control. Images are representative of 3 independent experiments. (**D**), His-tagged rPTH1R-ECD was incubated with recombinant catalytic domain of MMP14 at two enzyme/substrate ratios (rPTH1R-ECD only, 1:50, 1:10, and 1:10 with EDTA). The protein mixture were deglycosylated(left panel). Western blotting analyses used specific antibody indicated. Data are representative of three independent experiments. (**E**), Representative western blot trabecular bone protein lysate Mmp14+/+(left lane) and Mmp14-/-(right lane) mice at P14 (n=2). (**F**), Synthetic peptide from extracellular domain of PTH1R 55-67 was incubated with recombinant MMP14. The presence of 770Da suggests a fragment of VLQRPAS.

To test the hypothesis, we assayed the impact of co-expressing (Human influenza hemagglutinin) HA-tagged PTH1R with full-length MMP14 (MT1-FL) or catalytic inactive MMP14 (MT1-EA) in human 293T human embryonic kidney (HEK) cells (Fig. 3B). After deglycosylation with PNGaseF, Western blotting showed, in the presence of MT1-FL, a truncated form of PTH1R-HA with size close to 55 kDa is increased compared to empty vector and MT1-EA (Fig. 3B). To exclude that the cleavage is an artifact due to HA-tag insertion in the extracellular domain, a mouse *Pth1r* cDNA without HA-tag was reverse transcribed from extracted RNA and cloned into pcDNA3.1 expression vector (Fig. 3C). Like PTH1R-HA, a truncated protein fragment of PTH1R was detected when MT1-FL was expressed, confirming proteolytic processing of PTH1R exists both in mice and human (Fig. 3C). To exclude the possibility of indirect cleavage of PTH1R by MMP14 in 293T cells, a 6xHis-tagged recombinant extracellular domain of human PTH1R (PTH1R-ECD) was incubated with and without recombinant human MMP14(rhMMP14) overnight (Fig. 3D). Western blots showed two truncated fragments with size around 20 and 15kDa can be observed compared to intact PTH1R-ECD (Fig. 3D). Blocking MMP14 activity by EDTA inhibited PTH1R cleavage (Fig. 3D). To test if other MMPs/ADAMs participate in the cleavage of PTH1R, we assayed ADAM10, ADAM15 and ADAM17 and found the cleavage is restricted to MMP14 (Fig. S4A). Co-immunoprecipitation of PTH1R-HA and MT1-FL suggested that MMP14 can interact with PTH1R (Fig. S4B & C).

To test if MMP14 cleavage of PTH1R occurred *in vivo*, trabecular bone protein lysate was prepared from wild-type and *Mmp14^-/-^* mice at P14. A truncated 55kDa form of PTH1R can be detected in the *Mmp14^+/+^* trabecular bone lysates but not in those from *Mmp14^-/-^* mutants (Fig. 3E). However, an additional protein band at 110 kDa can be observed *in vivo* compared to secondary cell lines, possibly reflecting differences in posttranslational modification (48).The truncated 55kDa form of PTH1R was not present in lysates from cultured trabecular bone osteoblasts of *Mmp14^-/-^* mice (Fig. S4D), consistent with the role of MMP14 as a novel protease for PTH1R.

Given that rhMMP14 directly cleaves PTH1R-ECD into fragments with size around 20kDa and 15kDa, we propose at least one putative cleavage site exist around amino acid 55 to 65 in PTH1R. Computational prediction also suggest a possible cleavage site exist at amino acid around 61(49). To test this, we incubated rhMMP14 with a synthetic peptide with amino acid sequence 55-67 from PTH1R. Using mass spectrometry, we found a peak at 770 Da consistent with molecular mass of a VLQRPAS fragment, suggesting amino acid 61 is one of the cleavage sites of MMP14 (Fig. 3F).

### MMP14 inhibits PTH/PTH1R signaling

Prior molecular and structural studies have demonstrated that the exon 2 encoding region of PTH1R, which harbors the cleavage site by MMP14, is dispensable for both binding to PTH/Pthrp and receptor function(50–53). Since MMP14 has a wide range of substrates ranging from collagens to transmembrane ligands, we hypothesized that cleavage of MMP14 promoted the degradation of PTH1R. To explore the molecular consequences of MMP14 cleavage on PTH1R, 293T cells stably expressing PTH1R transfected with MT1-FL or MT1-EA were challenged with PTH and their responses examined by calculating the relative amounts of phospho-CREB (p-CREB) (Fig. 4A & B). As expected, lower relative amounts of p-CREB, p-ERK and cyclic-AMP (cAMP) were observed after PTH challenge in 293T cells expressing MT1-FL compared to control, suggesting MMP14 inhibits PTH signaling by facilitating receptor degradation (Fig. 4B & C). We isolated trabecular osteoblasts from *Mmp14^/--^* mice in order to ensure only MMP14 deficient cells were analysed. These MMP14 deficient primary trabecular osteoblasts showed increased p-CREB activity in response to PTH (Fig. 4D). We titrated the level of MT1-FL against the same amount of PTH1R and found increasing MT1-FL further promoted PTH1R degradation (Fig. S5A). Consistent with previous reports, MMP14 is upregulated in response to PTH treatment (Fig. S5B). Overall, these results identify cleavage of PTH1R by MMP14 amino acid 61as a novel mechanism for modulating GPCR stability and subsequently attenuated PTH signaling (Fig. 4E & S5).

**Figure 4.**
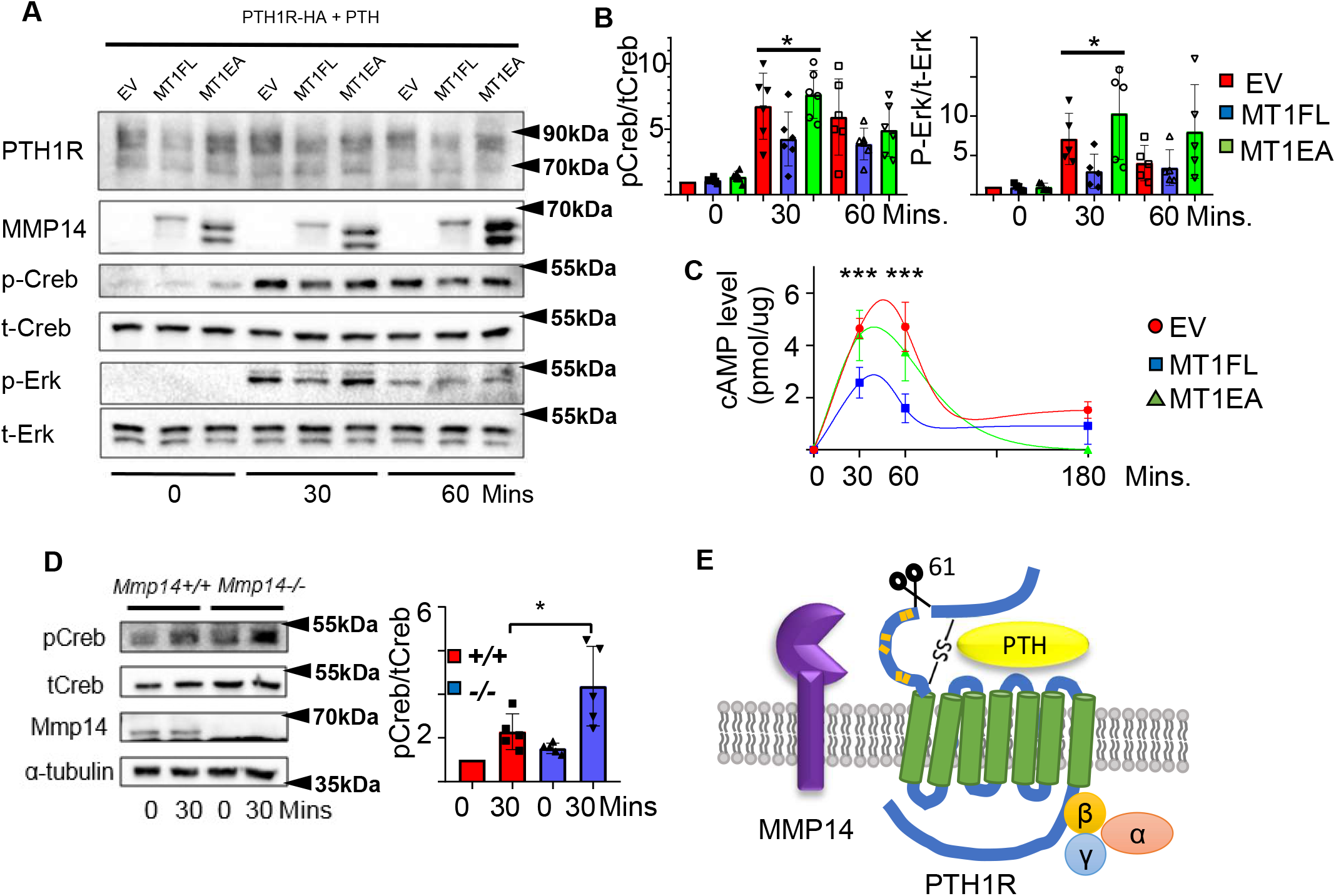
MMP14 inhibits PTH signaling. (**A**), MMP14 can inhibit PTH signaling in HEK293 cells. HEK293 cells stably expressing PTH1R-HA were transfected with empty vector(EV), MMP14(MT1-FL) and catalytic inactive MMP14(MT1-EA). Cells were challenged with 1×10-7M PTH in vitro for 0,30 and 60 minutes. PTH1R, MMP14, phospho-creb(p-creb), total-creb(t-creb), phosphor-Erk, total-Erk and loading control β -actin were analyzed by western blotting. (**B**), Relative level of p-creb/t-creb and p-Erk/t-Erk were quantitated and measured using ImageJ (n=6 for p-Creb/t-Creb and n=5 for p-Erk/t-Erk). One-way ANOVA was used to determine statistical significance. Data are presented as means ± SEM. ** p<0.01, * p<0.05. (**C**), HEK 293 cells expressing PTH1R were challenged with PTH for 30,60 and 180 minutes and the level of cAMP with MT1-FL,MT1-EA or empty vector were measured(n=5). One-way ANOVA was used to determine statistical significance. (**D**), Trabecular osteoblasts extracted from Mmp14+/+(red bar) and Mmp14-/-(blue bar) mice were challenged with 1×10-7M PTH(1–34). Cell lysate were analyzed by western blotting for p-Creb and t-Creb level (left). Right, quantification of relative level of p-Creb/t-Creb in trabecular osteoblasts extracted from Mmp14+/+ and Mmp14-/-mice (n=5). Data are presented as means ± SEM. ** p<0.01, * p<0.05, unpaired t test. (**E**), Schematic diagram showing MMP14 cleaves PTH1R at amino acid 61. Orange stripe indicates mutation sites. Seven-pass transmembrane domains are in green. PTH1R transduces signals via G-protein complex α,β,γ.

### HC-derived osteogenic cells respond to PTH which is enhanced in *Mmp14^ΔHC^*

Having demonstrated MMP14 can directly cleave PTH1R to negatively regulate its function, we next asked if PTH stimulates the osteogenesis of HC-derived cells and if MMP14 moderates PTH signaling in HC-derived cells *in vivo*. While treatment of mice with PTH inhibits apoptosis of osteoblast precursors and promotes their differentiation into mature osteoblasts (54), whether HC-derived osteoblasts respond and contribute to the anabolic response to PTH reported for osteoblasts is not known. To assess the contribution of HC-descendants to *Col1a1-*expressing mature osteoblasts induced by PTH treatment, the *C1-GFP* transgene was introduced into *C10Cre;RtdT* and *Mmp14^ΔHC^;RtdT* mice to label osteoblasts in bone (Fig. 5A). We examined the skeletal phenotype in *Mmp14^ΔHC^* mutants treated with PTH for 4 weeks (Fig. 5B). Compared to WT controls, PTH treatment in *Mmp14^ΔHC^* mutants caused a 26% and 23% greater increase in BMD and BV/TV, respectively, (Fig. 5B & C). PTH treatment dramatically increased the amount of HC progeny (tdT^+^, detected by RFP immunostaining) in *Mmp14^ΔHC^* trabecular bone (Fig. 5B, C).

**Figure 5.**
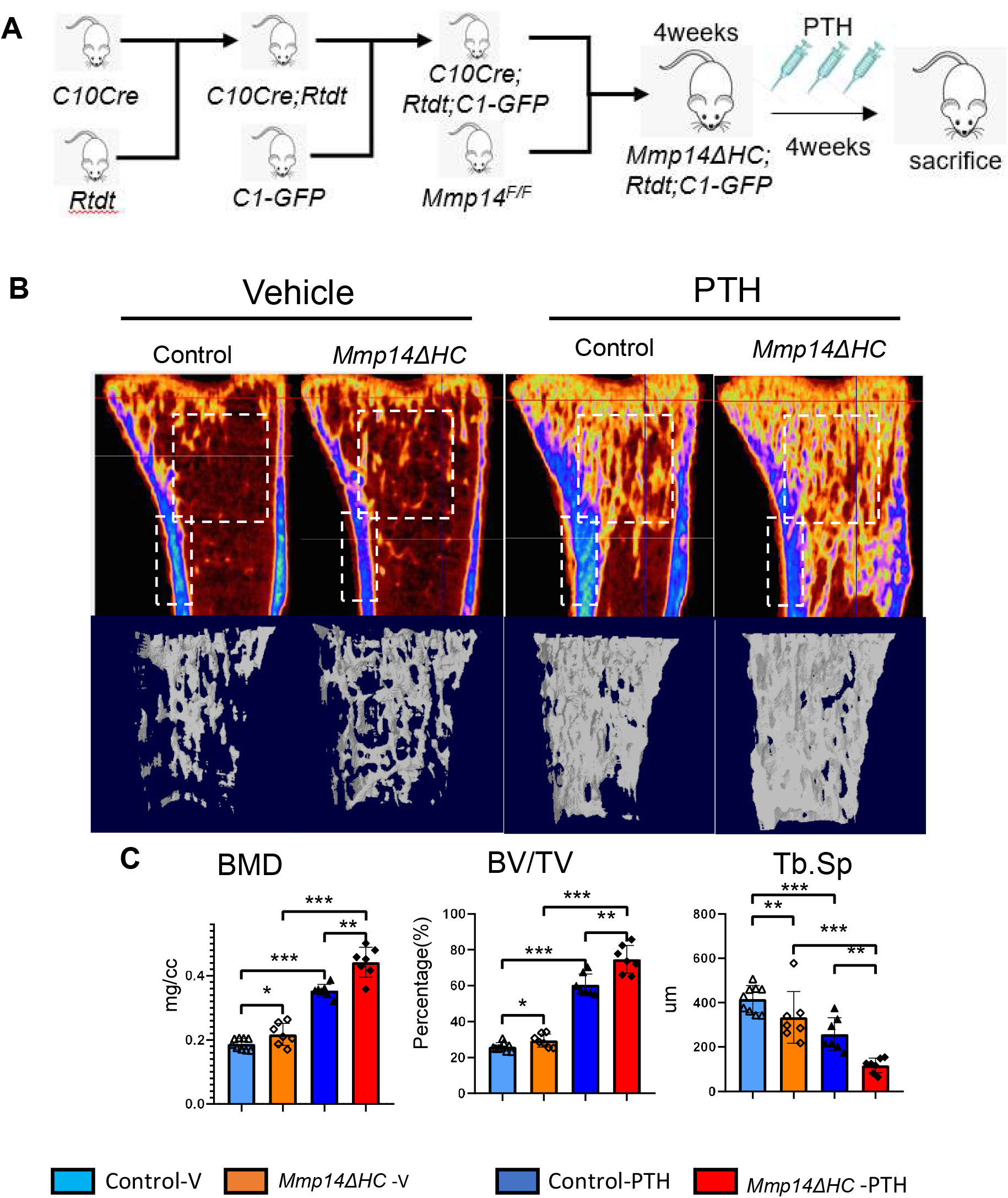
Increased anabolic response of HC-derived osteoblasts to PTH treatment in Mmp14ΔHC mice. (**A**), Schematic diagram showing experimental design for generation of Mmp14ΔHC; Rtdt;C1-GFP mice for Teriparatide treatment. (**B**), Sagittal and Reconstructed 3D image of Micro-CT of Control and Mmp14ΔHC mice treated with vehicle and PTH(1–34) for one month starting from P28 for 5 injections per week. (**C**), Statistical analysis of bone mineral density (BMD), bone volume over tissue volume ratio (BV/TV) and trabecular separation (Tra.Sp.) using Micro-CT in control (blue(n=9) and deep blue (n=7) bars) and Mmp14ΔHC (orange (n=7) and red (n=7) bars), mice treated with vehicle(blue and orange) and PTH(1–34)(deep blue and red), for one month. Data are presented as means ± SEM. ** p<0.01, * p<0.05, One-way ANOVA.

In compound *C10Cre;RtdT;C1-GFP* mice, mature osteoblasts, pre-osteoblasts and HC-descendants were labeled by GFP, OSTERIX and RFP, respectively. We found many more HC-descendants in the trabecular region that co-expressed OSTERIX, a marker of osteoprogenitors, and C1-GFP in response to PTH treatment than in controls (Fig. 6A, B). These HC-descendants showed positive pCREB^+^RFP^+^ staining confirming PTH signaling activity in HC-derivatives (Fig. 6C). HC-derived cells constituted approximately 50% of all trabecular osteoblasts with and without PTH treatment (Fig. 6D). HC-derived osteoblasts therefore represent a significant osteogenic population directly contributing to half of the total osteoblast population and capable of responding to PTH treatment in the same way as non-HC-derived osteoblasts (Fig. 6D). In *Mmp14^ΔHC^* mice, the number of RFP+ and RFP+GFP+ and RFP+OSX+ cells further increased compared to control mice treated with PTH, consistent with the heightened response to PTH treatment in *Mmp14* deficient HC-derived cells (Fig. 6D).

**Figure 6.**
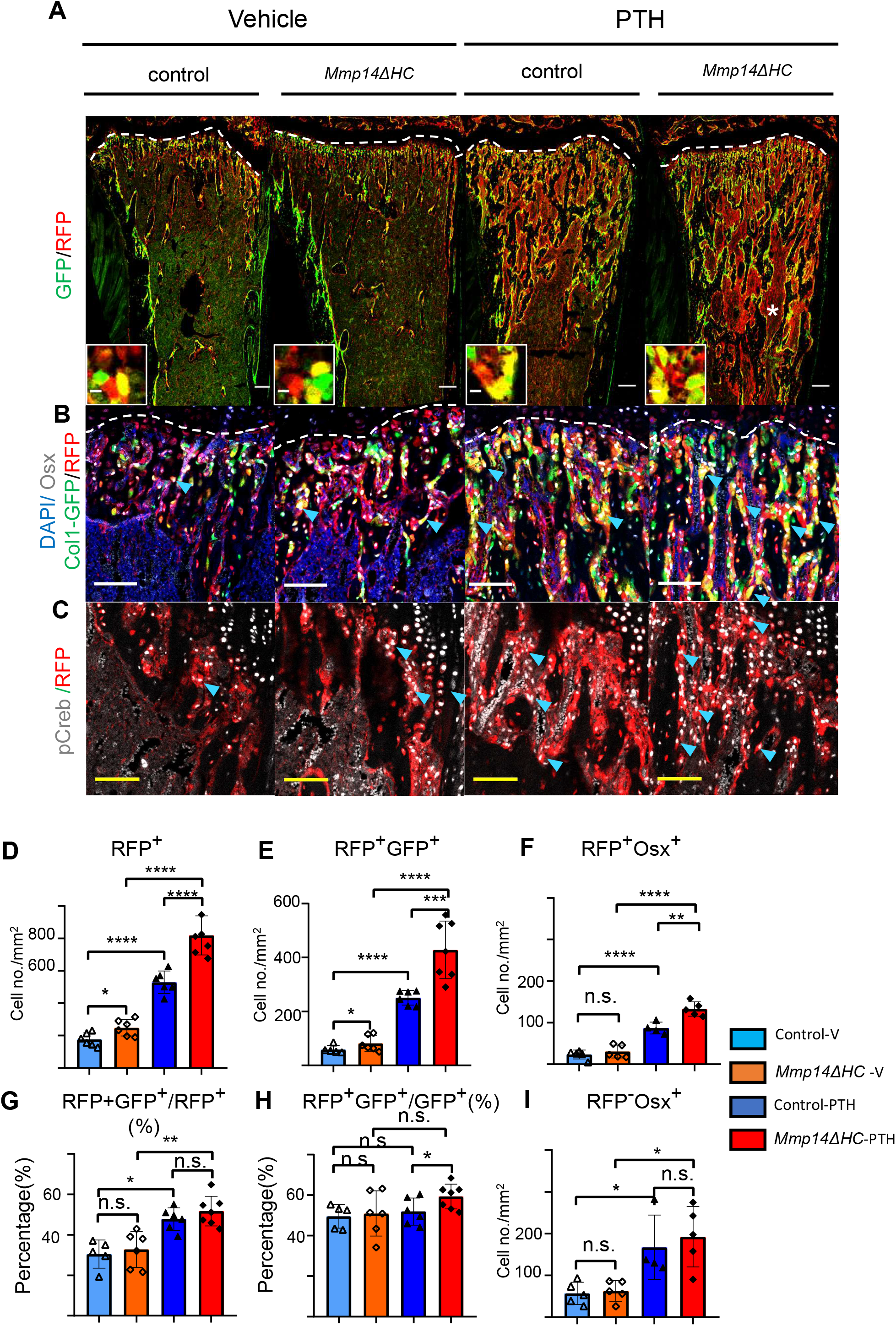
MMP14 controls expansion of chondrocyte descendants after PTH treatment. (**A**), Representative immunofluorescence staining of GFP (green) and RFP(red), marking HC-descendants and osteoblasts respectively. Mmp14ΔHC mutant mice (8 wk) and their littermate controls were treated with vehicle or PTH(1–34) for 4 weeks. Yellow dotted line demarcates chondro-osseous junction. Scale bar 50μm. (**B**), fluorescence colours mark nuclei, pre-osteoblasts and HC-descendants as for Fig 2. Scale bar 100μm. (**C**), Representative immunofluorescence staining of phospho-creb (pCreb, (grey) and RFP(red). Scale bar 100μm (**D-I**), Quantification of cell number /mm^2^: (**D**), RFP+ (HC-descendants); (**E**), RFP+GFP+ (mature HC-derived Osteoblasts); (**F**), RFP+Osx+ (HC-derived pre-osteoblasts), (**G**), ratio of RFP+GFP+/RFP+ cells, (**H**), ratio of RFP+GFP+/GFP+ cells and (**I**), number of RFP-Osx+ cells in Control(blue and deep blue) and *Mmp14ΔHC* (orange and red) mice (n=5). The data represent means ± SEM. *p* values were calculated using One-way ANOVA. Statistical significance of RFP+GFP+/GFP+ ratio is analyzed by students’ t-test(*= p < 0.05). The data points were pooled across multiple bone samples.

By contrast with enhanced osteogenesis in trabecular bone, BV/TV and cortical BMD were reduced in PTH-treated *Mmp14^ΔHC^* mutants compared to PTH-treated controls (Fig. S6A). There was no significant increase in RFP+GFP+ HC-derived endosteal cells in *Mmp14^ΔHC^* mutants. Although both HC-derived (RFP+GFP+) and non-HC derived (RFP-GFP+) endosteal cells responded to PTH treatment, the response to PTH was not enhanced in mutants, suggesting differences in the effect of MMP14 deficiency and PTH sensitivity of HC-derived endosteal cells compared to the non-HC-derived counterparts. Although the number of HC-derived RFP+ osteocytes were also not different between controls and mutants, there was a significantly enhanced response of HC-derived osteocytes to PTH in the cortical bone of *Mmp14^ΔHC^* mutants (Fig. S6A and B). These results may be relevant to the interesting observation that PTH treatment results in increased trabecular bone in patients but had no effect on endo-cortical, cortical or periosteal bone (55). Whether the discrepancy in outcome of PTH treatment or MMP14 deficiency on cortical bone and trabecular bone is related to a more transient effect on PTH signaling on bone synthesis in the former than in the latter and /or remodeling differences, are questions for future study.

### HC-descendants persist and contribute to the anabolic response to PTH in aged mice

Although, unlike in humans, the growth plate of mice does not close, in adult mice by one year of age, the hypertrophic cartilage is vestigial and endochondral ossification diminishes substantially (56). To test whether the HC-derived osteogenic cells could also contribute to the response to PTH treatment in older mice, we treated 1 year old *C10Cre;RtdT* mice with PTH for 4 weeks (Fig. 7A-C (i-iii)). Like in young mice, HC-descendants at 1 year old responded to PTH (1–34) treatment by producing more osteogenic progeny (Fig. 7C(i-iii)), suggesting sustained ability of HC-derived osteoblasts to contribute to PTH-stimulated osteogenesis with aging. Taken together our findings implicate a molecular link between MMPs and PTH signaling in regulating the activity of HC-derived osteoblasts in postnatal and adult mice (Fig. 8).

**Figure 7.**
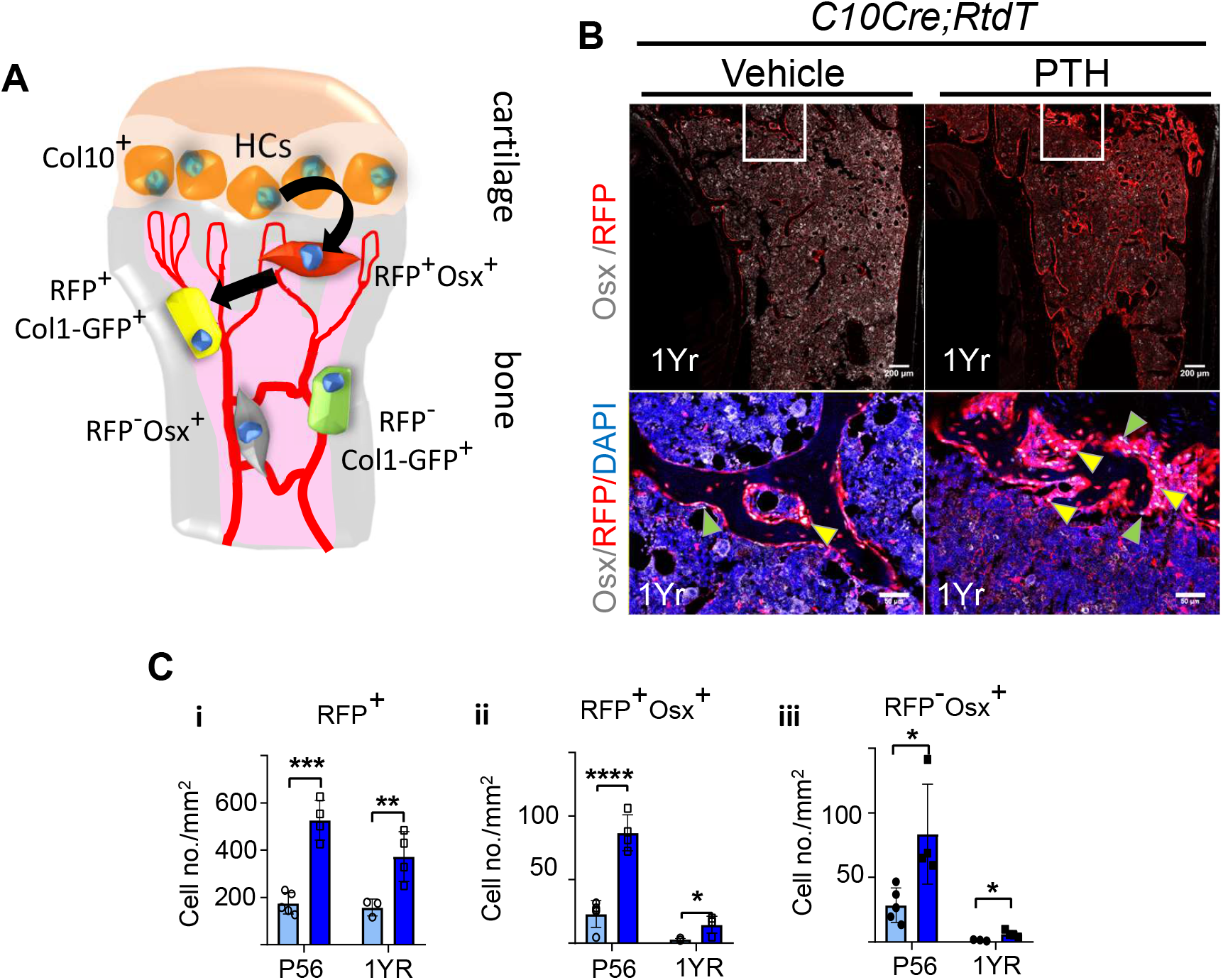
HC descendants persists in trabecular bone and contribute to PTH response in ages adult mice. (**A**), Schematic diagram showing ontogeny of HC derived and non HC-derived osteogenic cells and their markers. (**B**), Immunofluorescence staining of DAPI(blue), Osterix (grey) and RFP(red) for marking cell nuclei, pre-osteoblasts and HC-descendants respectively. Mice were treated with vehicle or PTH(1–34) for 4 weeks at five injections per week starting from 1 years old. Magnified picture of at the trabecular bone (white box) are also presented. Yellow arrows mark HC derived pre-osteoblasts. (**C**), P56 and 1 year mice treated with PTH: i-iii, quantification of number/mm2 of RFP+ cells, RFP+Osx+ cells, RFP-Osx+ cells. (n=3). Data are represented as mean ± S.E.M. p values were calculated using two-tailed unpaired t-test, * p<0.05, ** p<0.01, *** p<0.001. Data points were pooled measurements across multiple bone sections per animal.

**Figure 8.**
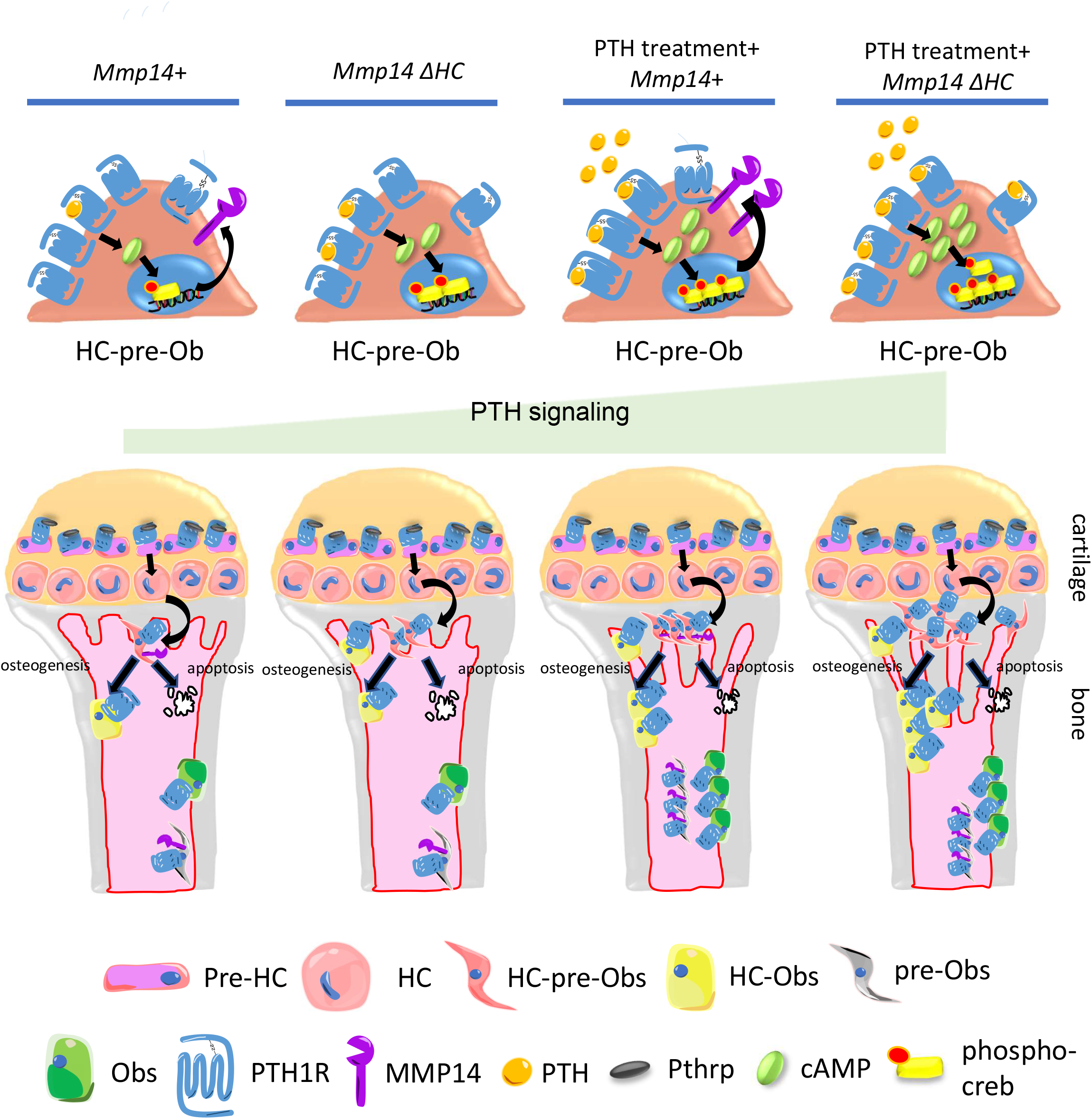
Model of MMP14 moderation of PTH signaling intensity and osteogenesis. Upon PTH binding, PTH1R generates cAMP signals to activate cAMP response element binding protein (Creb). Activation of PTH1R causes upregulation of MMP14. MMP14 in turn cleaves PTH1R to inhibit its signaling activity. In the chondrocyte lineage, pre-hypertrophic chondrocytes (pre-HC) differentiate to HCs, translocate to the subchondral space and subsequently to HC-pre-osteoblasts contributing to trabecular, endosteal and endocortical bone and can respond to PTH treatment. Together, MMP14 moderates the HC derivatives’ response to PTH ensuring a controlled supply of osteoblast precursors and thereby bone anabolism. Obs, osteoblasts, PHZ, Hypertrophic zone. HZ, Hypertrophic zone. TB, Trabecular bone.

## Discussion

Recent studies have established the concept that in the growth plate, resting chondrocytes function as skeletal stem cells for continuous supply of proliferating and pre-hypertrophic chondrocytes and HCs (12, 13). These cells form part of a continuum in which the HCs transition into the full osteogenic lineage, contributing to the formation of primary spongiosa and trabecular bone (9, 15, 17-19) and participating in bone regeneration and healing (7, 15, 17, 57). Here we show that a significant proportion of osteoblasts contributing to PTH-mediated bone formation in fact originated from chondrocytes. Hypertrophic chondrocytes serve as reservoir of progenitors suppling osteoblasts precursors and this process is controlled by MMP14 by direct cleavage of PTH1R. MMP14 inhibits PTH1R signaling and genetic targeting of MMP14 boosts PTH-mediated bone synthesis.

Our study has provided new in vivo insights into the properties and functional contribution of chondrocyte-derived osteoblasts to maintaining normal bone mass in response to PTH. The single-cell transcriptomic analyses and lineage tracing in mouse models show that HC-derived cells express genes characteristic of skeletal progenitors, mesenchymal cells, immature and mature osteoblasts, and terminally differentiated osteocytes even at maturity (P56) in trabecular and endosteal/endocortical bone. Despite this gross similarity there were a few genes that were differentially expressed between HC-derived and non-HC-derived osteoblasts, the functional significance of which is unknown and worthy of future study.

In our study we have found, by contrast to the decreased trabecular bone caused by global knockout of *Mmp14*, that conditionally inactivating MMP14 in the HC-derived lineage, results in increased bone by promoting their survival. Our findings identify MMP14 as a key protease that regulates the contribution of HC-derivatives to trabecular bone. MMP14 inhibits osteoclastogenesis via controlled release of soluble RANKL (46). RANKL has been reported to be a downstream target of PTH and a substrate of MMP14, which implies that MMP14 is downstream of PTH signaling, but the exact molecular relationship is unclear. Notably osteoclastogenesis was normal in the *Mmp14^ΔHC^* mutants, suggesting that the increased bone mass is intrinsic to HC-derived osteoblasts and unrelated to an impact on the production of soluble RANKL in the absence of MMP14.

Because inactivating MMP14 mediated intracellular signaling constitutively by mutating the cytoplasmic tail of MMP14, leads to increased trabecular bone and decreased adipocytes in mice, it has been suggested MMP14 regulates the osteogenic versus adipogenic cell fate decision of differentiating bone marrow skeletal stem cells (38). However, there was no effect on the frequency of HC-derived adipocytes in *Mmp14^ΔHC^* mutants suggesting the increased osteogenesis is not caused by a change in HC cell fate.

GPCR is the largest family of transmembrane proteins with contributing to ∼35% of encoded proteins. Around 3-40% of drugs are derived to target GPCRs. Few preliminary reports propose proteolytic modification of GPCRs by MMPs or ADAMs might modulate their signaling activity (48, 58). A recent paper proposed that cleavage of PTH1R could regulate its downstream signaling (59). Our study has uncovered that PTH1R, a class-B G-protein coupled receptor, is a novel direct substrate for MMP14 which, MMP14, by cleaving PTH1R acts to inhibit PTH1R signaling, The detailed structure of PTH1R and the lack of functional impact of HA-tagging in this domain suggests the unstructured region encoded by exon 2 does not participate in receptor-ligand binding nor signal transduction (52, 53, 60). Our results demonstrate proteolytic modification of this region directly influences GPCR signaling via GPCR stability. We propose a novel paradigm of MMP14 activity in which it also mediates direct titration of PTH signaling in the HC-osteoblast lineage. In this model the intensity of PTH stimulation of osteogenesis is curbed via MMP14 cleavage of the ectodomain of PTH1R. Hence, inactivation of MMP14 in HC-descendants spares the ectodomain of PTH1R from cleavage, resulting in an increased anabolic response to PTH and therefore places MMP14 also upstream of PTH signaling. Since MMP14 is upregulated by PTH signaling in osteogenic cells *in vivo* (46), it is possible MMP14 is also involved in a negative feedback protection mechanism to constrain osteogenesis in response to PTH challenge.

Maintaining appropriate amount of bone throughout life, mediated by hormones such as PTH, is critical for healthy aging and quality of life. However, there is a gap in knowledge on the specific contributions of individual cell types to maintaining homeostasis in bone because of the myriad of signaling pathways and diverse cell-type responses. The importance and need of a molecular and functional definition of the contribution of specific cell types in bone is illustrated by the long-term systemic treatment of osteoporosis by Teriparatide (PTH 1-34). The treatment is complicated by side effects because of the physiological and pleiotropic effects of PTH signaling on different cells in the whole body. We have shown that HC-derived osteoblasts contribute to the anabolic activity in bone, responding to PTH in the same way as their non-HC-derived counterparts, and constitute a significant proportion of the osteogenic precursors stimulated by Teriparatide (PTH 1-34) treatment, even in adult mice. This response to PTH occurs in early postnatal and mature mice (1 year). Questions for future study are whether the response is as robust in aged mice (2 years or older) and if, compared to other sources of osteoblasts, the history of PTHRP/PTH1R signaling in pre-hypertrophic chondrocytes (12) produces a molecular memory that is advantageous for the ability of HC-descendants to mount a response to PTH. Whether a defect in the HC-lineage progression underlies the increase in trabecular bone in chondrocyte-mediated *Mmp13* deficiency (28) and the exact underlying molecular mechanism(s) are questions that should be followed up. It would also be important to explore whether there is a causal link between the chondrocyte-osteoblast lineage and human bone-wasting disorders.

Our study illuminates understanding of the relative contribution of different cell types to maintaining bone homeostasis and highlights the importance of HCs as a source of pre-osteoblasts capable of rapid response to PTH treatment. The minor differences in transcriptomic signature between HC-derived and non-HC-derived osteoblasts and their co-expression of *Mmp14* and *Pth1R,* suggest that the same paradigm of MMP14 mediated titration of PTH signaling applies in all osteoblasts. Addressing this possibility purely in non-HC-derived osteoblasts is hampered by the lack of appropriate cre reagents that can target osteoblasts in the perichondrium specifically and the intrinsic skeletal defects in Osterix–cre mice (61). The discovery that MMP14 titrates the response to PTH and bone anabolism in vivo, contributes new insights into the mechanism of titrating PTH control of osteogenesis and bone homeostasis and has profound significance for understanding bone metabolism with implications for the development of treatments for bone wasting diseases such as osteoporosis and in overcoming the side effects of Teriparatide (PTH 1-34). Our work also contributes to understanding the scientific basis of PTH therapy and provides molecular insight into potential enhancement of treating skeletal diseases. It has been suggested that a therapeutic approach could be to reduce bone resorption by targeting the myeloid lineage which would require inactivating both MMP9 and MMP14 (43). By contrast a pan osteoblast-specific or HC-derived osteoblast-specific targeted manipulation approach would have a significant advantage as MMP14 activity (and therefore PTH signaling) only needs to be manipulated in a single cell-type and it avoids systemic intermittent injection of PTH which would thereby ameliorate side effects.

## Materials and Methods

### Genetically modified mouse strains

*Mmp14^-/-^* and *Mmp14* ^flox/flox^ mice were generated as described previously (29, 35). *Col10a1-Cre* and *Col10a1-CreERT* mouse lines are described in Yang et.al (15). *Col1a1-2.3-GFP* and *Rosa26* reporter transgenic mice were obtained as previously described (62, 63). PTH delivery to mice was performed as described in Balani et.al (54). In brief, PTH(1–34) (acetate, Bachem) was dissolved in 0.01M acetic acid, 150 nM NaCl and subcutaneously injected to mice at 400 ng/g body weight for five days per week (54). All animal experiments were approved by the Institutional Animal Care and Use Committee of The University of Hong Kong and performed in accordance with guideline of the Committee on the Use of Live Animals for Teaching and Research of The University of Hong Kong.

### Microcomputed tomography (MicroCT) imaging

The tibia and femora were collected and fixed in 100% absolute ethanol overnight at 4 °C. The tibia was sent to scanning by the eXplore Locus SP System (GE company, U.K.). Acquired images were processed using the Dataviewer, CTAnalyser and CTVolume from Bruker, U.S.A. Morphometric values including bone mineral density (BMD), bone volume/ tissue volume (BV/TV), trabecular thickness (Tb.Th), trabecular separation (Tb.Sp) and trabecular number (Tb.N) are presented. Samples from Mmp14F/F;C10Cre and Mmp14F/-;C10Cre were pooled for MicroCT analysis.

### Histology, immunostaining and *in situ* hybridization

Hematoxylin and Eosin(H&E) staining, immunostaining was performed as previously described (15). Briefly, mouse tibiae were fixed in 4% (weight/volume) paraformaldehyde overnight at 4 °C with agitation. Tissues were decalcified in 10% EDTA for three washes for three days. The samples were further dehydrated and embedded in wax. Immunostaining and *in situ* hybridization were performed as previously described. Samples from *Mmp14^-/-^*, *Mmp14^+/+^*, *Mmp14^F/F^;C10Cre*, *Mmp14^F/-;^C10Cre* and *Mmp14^F/+^;C10Cre* were analyzed.

### TUNEL labeling and Edu incorporation assay

Proliferation and apoptosis assays were described in Yang et.al. (15). Edu labeling was performed with intraperitoneal injections of Edu at 150 mg/kg body weight and the mice were sacrificed 2 hours later. Labeled cells in paraffin sections were detected using Click-iTR EdU imaging kits from Invitrogen. *in situ* terminal deoxynucleotidyl transferase deoxyuridin triphosphate nick end labeling (TUNEL) assay was performed using *in situ* cell death kit from Roche (Basel, Switzerland).

### Quantitation of cells

Fluorescence images were processed with ImageJ (developed by National Institutes of Health, USA). For young mice at P3 or P10, the number of cells was counted manually. For mice aged 2 months, images were taken using Carl Zeiss LSM confocal 800 system (Carl Zeiss, Oberkochen, Germany). Overlap of DAPI with RFP (tdTomato), GFP or OSTERIX was considered as one single cell as in Fig.6. Three separate bone sections from each animal were counted blindly in a region below the chondro-osseous junction. Cells were quantitated using ImageJ software and averaged by a specified area 4mm below chondro-osseous junction for 2 Months old mice. Statistical analyses were performed using Graphad Prism 8.0. Predetermined sample sizes were not used in statistical methods. Unpaired two-tailed student’s t-test was used. p< 0.05 was considered statistically significant.

For quantification of vascularized area below the chondro-osseous junction, blood vessels are stained with the endothelial cell marker ENDOMUCIN (EMCN). Vascularized area below the chondro-osseous junction is measured by manual drawing of blood vessel area circled by ECMN staining with ImageJ. Measured area is represented as mm^2^. Quantification of osteocytes were identified by their location in cortical bone matrix.

### X-gal staining

β-galactosidase staining was performed as previously described(15). Briefly, Harvested embryos were prefixed in 4% PFA for 1 hour at 4 °C. Prefixed samples were washed in washing buffer 3 times and were stained in X-gal solution overnight.

### Primary osteoblast culture

Extraction of trabecular osteoblasts was described in Chan et.al. (34). Tibiae and femora and the cartilage were carefully removed at P14. The entire tibia and femur were subjected to centrifugation to remove the bone marrow. Cortical bone was also removed and the remaining trabecular were subjected to serial digestion. Cells released after the third and fourth digestion were collected and the tdTomato-positive cells were clearly seen under fluorescent microscopy. The HC-descendants were first cultured in alpha-Minimum Essential Medium (αMEM) supplemented with 50 μg/ml penicillin/streptomycin, 12.5 μg/ml gentamycin and 0.5 μg/ml fungizone for three days and were subsequently differentiated in αMEM containing 50 mg/ml ascorbic acid, 10mM β-glycerophosphate, 10 μg/ml collagen type I (Stemcell Technologies, Catalog # 07001), 50 μg/ml penicillin/streptomycin, 12.5 μg/ml gentamycin and 0.5 μg/ml fungizone for seven days. Matrix producing osteoblasts (identified by *C1-GFP* fluorescence) were then challenged with 1x 10^-7^M PTH(1–34) and collected at indicated time points. Cells were washed with ice-cold PBS and lysed using Radioimmunoprecipitation assay (RIPA) buffer [150 mM NaCl, 0.5% sodium deoxycholate, 0.1% SDS (sodium dodecyl sulphate), 50 mM Tris-HCl, pH 8.0 supplemented with cOmplete™, EDTA-free Protease Inhibitor Cocktail (Roche, cat no: 11873580001)]. Western blotting was performed as previously described (34).

### Transfection and cell experiments

For testing *in vitro* cleavage of PTH1R by MMP14, HEK293 cells were seeded on 12 or 6 well dishes at 30% confluence and transfected with vectors(pcDNA3.1) expressing MT1-FL (Full length MMP14), MT1-EA (inactive MMP14) and PTH1R at 60-70% confluence with Fugene HD (Promega) according to manufacturer’s instructions. Cells were collected 48-72 hours after transfection. To test for cellular response to PTH, stable cell line expressing PTH1R was prepared by transfecting 293 cells with pcDNA3.1-PTH1R and was selected using G418 according to manufacturer’s instructions. Stable clones expressing PTH1R were examined by western blotting. To test for response to PTH, 1×10^-7^M PTH was added 48-72 hours after transfection with MT1-EA or MT1-FL and cell was collected at corresponding time points.

### cAMP ELISA assay

Cyclic adenosine monophosphate (cAMP) was measured using cAMP complete ELISA kit from Enzo Life Sciences (Cat. No: ADI-900-163). 293T Cells expressing PTH1R were transfected with empty vector, *Mmp14* and *Mmp14-EA*. After 48-72 hours, the cells were challenged with 1×10^-7^ M PTH (1–34) and subsequently lysed in 0.1M HCl in 0, 30 and 60 minute time series. The supernatant was collected and assayed according to manufacturer’s instructions.

### Selection of PTH1R expressing stable clone

To establish a 293 cell line which stably expresses PTH1R, a pcDNA3 vector from Invitrogen containing a neomycin selection gene, a human PTH1R coding sequence with a modified HA-tag at exon2 provided by Prof. Martin J. Lohse and a ORF human PTH1R (Genscript, cat.no. OHu15045D) was transfected into 293 cells. After 2 days, the transfected cells were selected with G418 at 1mg/ml for approximately 1 week. Individual clones was picked by pipetting and screened for PTH1R expression by western blot.

### Extraction of protein from mouse trabecular bone

Trabecular bones were harvested from the *Mmp14^-/-^* and littermate control mice, followed by homogenizing bones in RIPA with a bullet blender for 15 minutes at 4 ℃. After leaving on ice for 10 minutes, the samples were centrifuged at 13, 000 × g for 15 min at 4 ℃. Next, the supernatants were collected and processed for Bio-Rad protein assay for protein concentration determination.

### *In vitro* cleavage of rPTH1R and peptide fragments from PTH1R

Peptide fragments covering amino acids 55-66 (Genscript) and His-tagged recombinant extracellular domain of Human PTH1R (Cat: 12232-H08H, Sino biological) were used. The peptides and recombinant proteins were incubated with recombinant MMP14 (BML-SE259-0010, Enzo Lifesciences) for 16 hours and the mixture was analyzed using western blotting or MALDI-TOF mass spectrometry (Centre for Proteomics, The University of Hong Kong).

### Western blot analysis

Western blotting was performed as described previously (34). Briefly, RIPA cell lysis buffer was prepared using 150 mM NaCl, 1% NP-40, 50 mM Tris, 0.1% SDS and 0.1 % sodium deoxycholate. Protein lysate was then blotting on a polyvinylidene fluoride (PVDF) membrane (Millipore, MA, USA) and transferred for 1 hour at 15V using Trans-Blot® SD Semi-Dry Transfer Cell (BioRad). The PVDF membrane was blocked with 5% milk in Tris-buffered saline with 0.1% tween-20 (TBST) for 1 hour and incubated with agitation with primary antibody overnight at 4°C. The membrane was further incubated in secondary antibody conjugated to HRP diluted in blocking buffer for 1 hour at room temperature with agitation. Non-specific binding of HRP-secondary antibody to the PVDF membrane was washed for three times with TBST each for 3-5 minutes. To develop the signals for chemiluminescence of the targeted proteins, WesternBright ECL kit (K-12045-D50) from Advansta was used according to manufacturer’s instructions. After incubation with the ECL substrate, the PVDF membrane was taken to the ChemiDocTM MP Imaging system for the visualization of target proteins.

To detect the cleaved fragments of PTH1R by western blotting, the cell lysate was either not boiled or heated to 70℃ for 5 minutes for denaturation. Boiling will cause PTH1R containing 7-transmembrane domain to aggregate. EDTA was added to the RIPA cell lysis buffer to a concentration of 5mM to inhibit membrane metalloproteinase activity. PTH1R was deglycosylated by 1hour (secondary cell lines) or overnight incubation (primary cells or tissue lysate) with PNGaseF at 37°C (NEB, cat no, P0704S). 1µl of 1M DTT was added to 50µl of cell lysate to ensure complete dissociation of disulphide bonds linking cleaved extracellular domain of PTH1R. The processed protein fragments were analyzed by SDS-PAGE.

### Co-immunoprecipitation assay

Co-immunoprecipitation assay was performed as previously described(34). Briefly, *in vitro* cultured cells were transfected with PTH1R-HA and MT1-FL expression plasmids. After 48-72 hours, cells lysed using ice-cold IP lysis buffer (pH 7.5, 50 mM Tris-HCl, 150 mM NaCl, 1% NP-40, 0.5% sodium deoxycholate, 1 tablet protein inhibitor cocktail, 5mM EDTA) containing proteinase inhibitor cocktail. The cell lysate was centrifuged at 13,000 × g for 10 min at 4 °C, the supernatant was cleared with 50µl of Protein agarose G (Roche cat no. 11719416001) for 3 hours at 4 °C and subsequently incubated with 50 µl of Protein agarose G with corresponding primary antibody overnight at 4°C. The protein agarose complex was pelleted, resuspended with IP buffer two times, followed by washing in high salt (pH 7.5 50mM Tris-HCl, 500mM NaCl, 0.1% NP40, 0.05% sodium deoxycholate) and low salt(pH 7.5 50mM Tris-HCl, 0.1% NP40, 0.05% sodium deoxycholate) IP buffer. 30 μl 1× loading buffer was used to the antibody and G-agarose complex and heated to 65 °C for 10 min, followed by SDS-PAGE and western blotting.

### TRAP staining

Osteoclasts were detected by Tartrate resistant phosphatase (TRAP) staining. After rehydration of sections, they are incubated with Napthol-Ether Substrate at 37℃ for 1 hour. After that, they are changed directly to Pararosaniline solution freshly made by mixing 1:1 ratio of sodium nitrile solution and Pararosaniline Dye for color development. TRAP-positive, multi-nucleated (> 3 nuclei) osteoclasts are stained by red color and the section is counterstained by hematoxylin for contrast.

### Isolation of cells for single-cell transcriptomics

Single cells from the growth plate and trabecular bone were harvested, disassociated, digested, and sequenced as follows. For neonatal (P6) mice, cells were isolated from whole tibia (n=2) including the growth plate (except secondary ossification centre at both ends) of a *C10Cre;Irx3^+/ΔHC^Irx5^+/-^;R26^td/+^* heterozygote previously shown to have normal bone mass (19). The tibiae (n=2) were diced into fragments and digested with digestion solution (0.25% dispase, 0.25% collagenase type II in HBSS (Sigma-Aldrich, Cat. No: H6648)) (5ml in total) at 37 °C on a shaker for 1 hour. Similarly, the trabecular bone from tibiae (n=2) from an adult (P56) mouse (*Mmp14^ΔHC/+^;R26^td/+^*) were harvested. Osteogenic cells were isolated by a modification of protocol described by Chan et al (34). In brief, the first two suspensions obtained after 20 minutes sequential digestions, containing most of the blood cells, were removed, and fresh digestion solution was added. The single cell suspension enriched in Tdtomato+ cells (>10% Tdtomato+ by microscopy) released by the third digestion, was passed through a 40 µm filter. The cells were pelleted by centrifugation and washed twice with Hanks’ Balanced Salt solution (HBSS, Cat. No H6648, Sigma-Aldrich), resuspended in HBSS and the single cells encapsulated for library preparation and sequencing using the Chromium single cell platform (10X Genomics Inc.) at The University of Hong Kong, Centre for PanorOmic Sciences (HKU CPOS) per manufacturer’s protocol. Library size and concentration were determined by Qubit, quantitative PCR and Bioanalyzer assays. Viabilities of 69% and 74.5% were recorded for the P6 and P56 samples, respectively. A raw total input of 22750 cells was estimated for each sample.

### Single-cell RNA sequencing and bioinformatics analyses

Library preparation and Illumina (Novaseq 6000) sequencing (151bp paired end) were performed at HKU CPOS, at 150 Gbp throughput per sample. In both samples, 70% of bases had quality scores ≥ Q30. Cells with high percentage mitochondrial RNA and ribosomal RNA were filtered out for quality control. The raw data were aligned to mouse genome (mm10), tdTomato sequence (1431bp) and the Woodchuck Hepatitis Virus Posttranscriptional Regulatory Element (WPRE) sequence (593 bp), using Cell Ranger (v3.1.0). The R26 tdTomato cre-reporter vector contains the WPRE sequence downstream of the tdTomato sequence to reduce readthrough transcription (26). Since the current single-cell technology generates sequences ∼500bp upstream from the 3’ end of transcripts, tdTomato sequences may not always be detected due to the addition of WPRE. We confirmed that tdTomato and WPRE are linearly correlated but with enhanced detectability in the latter (Fig. S1 C and D). As such, we used WPRE as a surrogate for tdTomato. We assigned a cutoff value of WPRE>14 to match our experimental determination of the frequency of tdTomato-positive HCs ((15, 19)∼75%), which also corresponds to a saddle point in the WPRE distribution (Fig. S1 B-E). This cutoff was used to separate the osteogenic cells into HC-derived osteoblasts and non-HC-derived osteoblasts (Fig. 1 C, K, N). A kernel logistic regression model was trained to predict the boundaries between HC-derived osteoblasts and non-HC-derived osteoblasts, using the KLR package in R (64) (Fig. S1F). To isolate endochondral cells, cells with fewer than 1,600 genes or clusters expressing Ptprc (CD45) were excluded. Data were further processed with Seurat (v3.9.9)(65). In all, 3,570 (average 3,712 genes per cell) and 126 (average 1,103 genes per cell) endochondral cells were found in the P6 and P56, respectively. Population signatures were identified by comparing every population against all other cells, using Seurat’s FindAllMarkers function, with an FDR cutoff of <0.05, log_2_(fold-change)>1, and expressed percentage point difference >25%. From these signatures, eight endochondral bone populations were identified in P6 based on the literature (Fig. 1H), including chondrocytes, pre-hypertrophic chondrocytes, HCs, proliferating chondrocytes, immature osteoblasts, mature osteoblasts (5, 23, 24, 66, 67). In P56, only two osteogenic clusters were identified, immature and mature osteoblasts. The osteogenic cells of P6 and P56 were integrated using the canonical correlation analysis (CCA) approach (Fig. 1J). Gene-ontology analyses were performed using GSEA (gsea-msigdb.org).

### Statistics

Results were represented as mean ± SD. Statistical evaluation was done by nonparametric 2-tailed Student’s *t* test using GraphPad Prism version 8 for Microsoft (www.graphpad.com). ONE-WAY ANOVA for multiple testing was used for Fig. 4, Fig. 5. *P* < 0.05 was considered significant.

## Acknowledgments

We thank Henry Kronenberg for sharing mouse reagents and for critical input and advice on the study; Danny Chan, and Reinhard Faessler for helpful discussion, and Martin Loshtse for providing the PTH1R-HA expression vector.

## Author Contributions

Conceptualization: TLC, KSEC.

Methodology: TLC, MK, ZT, KYT, AXY.

Data analyses and interpretation: TLC, PC, MK, AXY, KSEC.

Figures: TLC, PC, AXY.

Supervision: KSEC, ZZ.

Writing—original drafts: TLC, KSEC

Writing—review & editing: TLC, KSEC, ZZ, AXY.

Funding acquisition: KSEC.

---

**Supplementary Figure 1.**
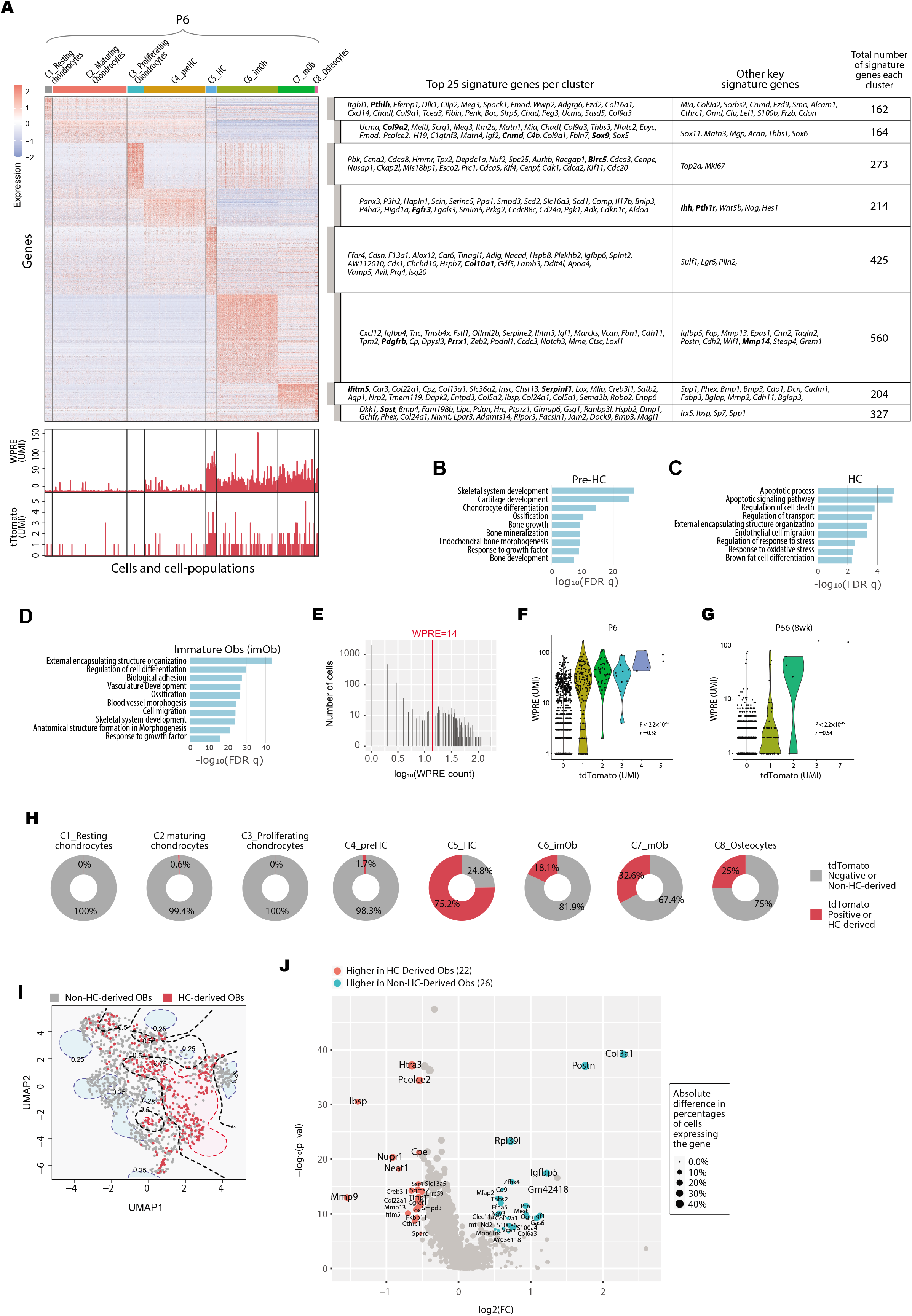
Integrative analyses of endochondral bone sub-populations at P6 and P56. (A), heatmap showing the signature genes of the 8 cell clusters in the P6 sample. Both the top 25 signature and certain key signature genes were listed. WPRE: Woodchuck Hepatitis Virus (WHV) Posttranscriptional Regulatory Element. (B-D) Gene Ontology analysis of significant cellular processes involved in Pre-HCs(B), HCs(C) and Immature-Obs(D). (E), A histogram showing the expression level of tdTomato and WPRE per cell. An optimal cutoff at WPRE>14, above which a cell was considered HC-derived, was chosen such that it corresponds to the saddle point of the histogram. (F-G),Violin plots showing the correlation between expression levels of WPRE and tdTomato in P6 (C) and P56 (D) samples. (H), Piecharts showing the percentages of tdTomato positive cells in each of the 8 clusters in P6. For cells in the C6, C7 and C8, these will correspond to being HC-derived. (I), Scatter-plot showing HC-derived Osteoblasts and non-HC-derived Osteoblasts. Contours are the boundaries as predicted by a kernel logistic regression model on the descendants vs non-descendants. Red contours represent descendant regions. Blue contours represent non-descendant regions. Black curves represent the mid-line between descendants and non-descendants. (J), Volcano plot showing the DEGs between HC-derived and non-HC-derived Osteoblasts in the integrated data of P6 and P56. preHC: pre-hypertrophic chondrocytes; HC: hypertrophic chondrocytes; imOb: immature osteoblasts; mOb: mature osteoblasts.

**Supplementary Figure 2.**
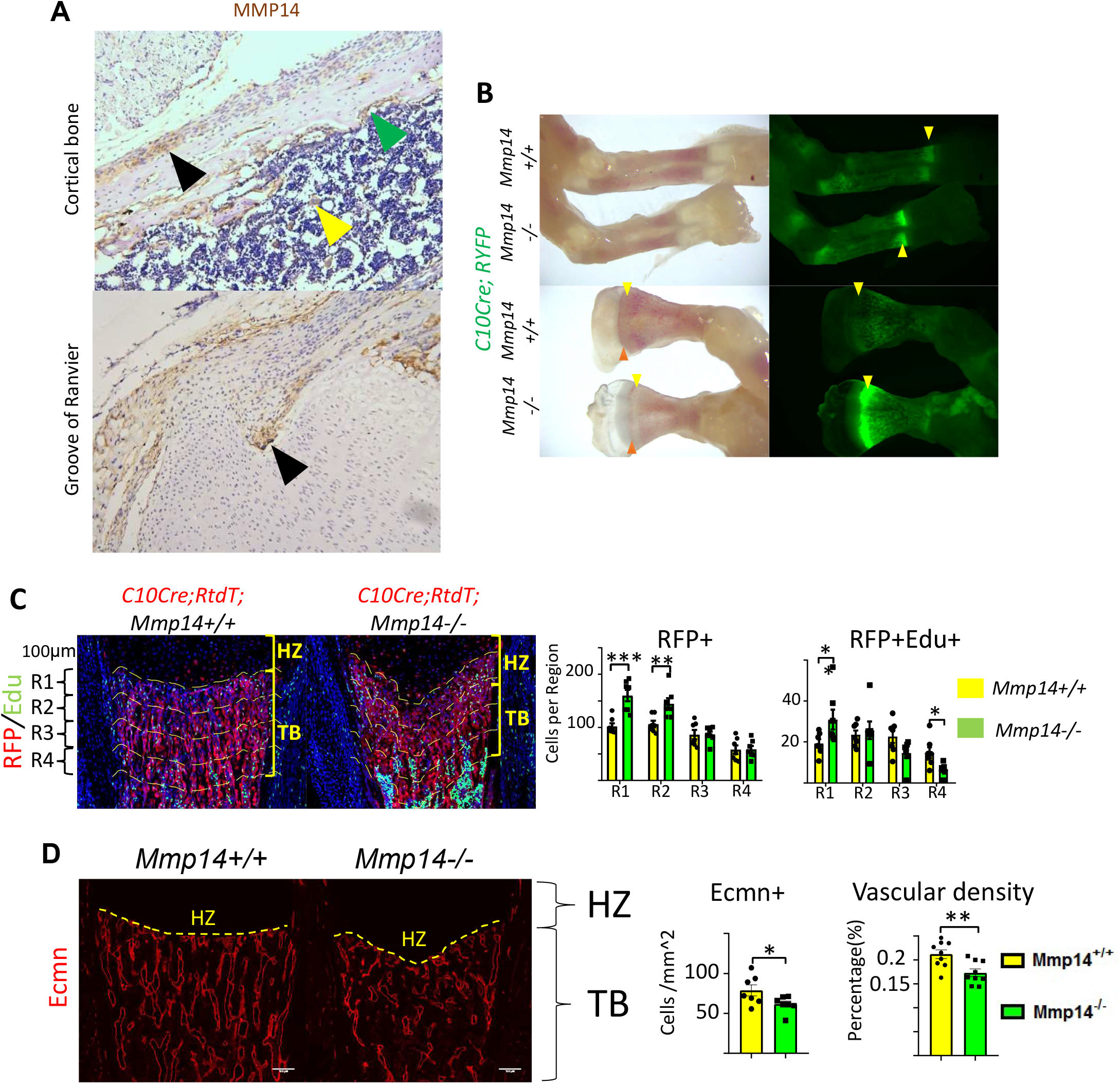
Abnormal localization of HC-descendants in *Mmp14^-/-^* mice. (A), Cells in cortical bone and groove of ranvier at P10. Yellow arrow indicates Megakaryocytes. Green arrow indicate endosteal bone lining cells. Black arrow indicates perichondral cells. (B), Bright field and fluorescence images of radius, ulna and scapula of *C10Cre;RYFP;Mmp14^+/+^* and *C10Cre;RYFP;Mmp14^-/-^* mice at P3. Green fluorescence label HC-derived cells. Red arrows mark chondro-osseous junction. Yellow arrow marks trabecular region. HZ represents hypertrophic zone. (C), Immunofluorescence staining of RFP and Edu-labeled cells counterstained with DAPI in *C10Cre;RtdT;Mmp14+/+* and *C10Cre;RtdT;Mmp14^-/-^* tibia at P3. For quantification, the trabecular bone is divided into four zones, each 0.1 mm in thickness. The number of RFP+(red), RFP+Edu+(red and green) cells in each region were counted. Data are presented as means ± SEM ** p<0.01, * p<0.05, unpaired *t* test. (D), Immunostaining of Endomucin (Endo) labeling blood vessels below chondro-osseous junction comparing wild-type (yellow bar)and Mmp14-/-(green bar) mice at P3. The vascular density and number of Ecmn cells are presented (n=9). Data are presented as means ± SEM ** p<0.01, * p<0.05, unpaired t test. HZ, hypertrophic zone. TB, trabecular bone.

**Supplementary Figure 3.**
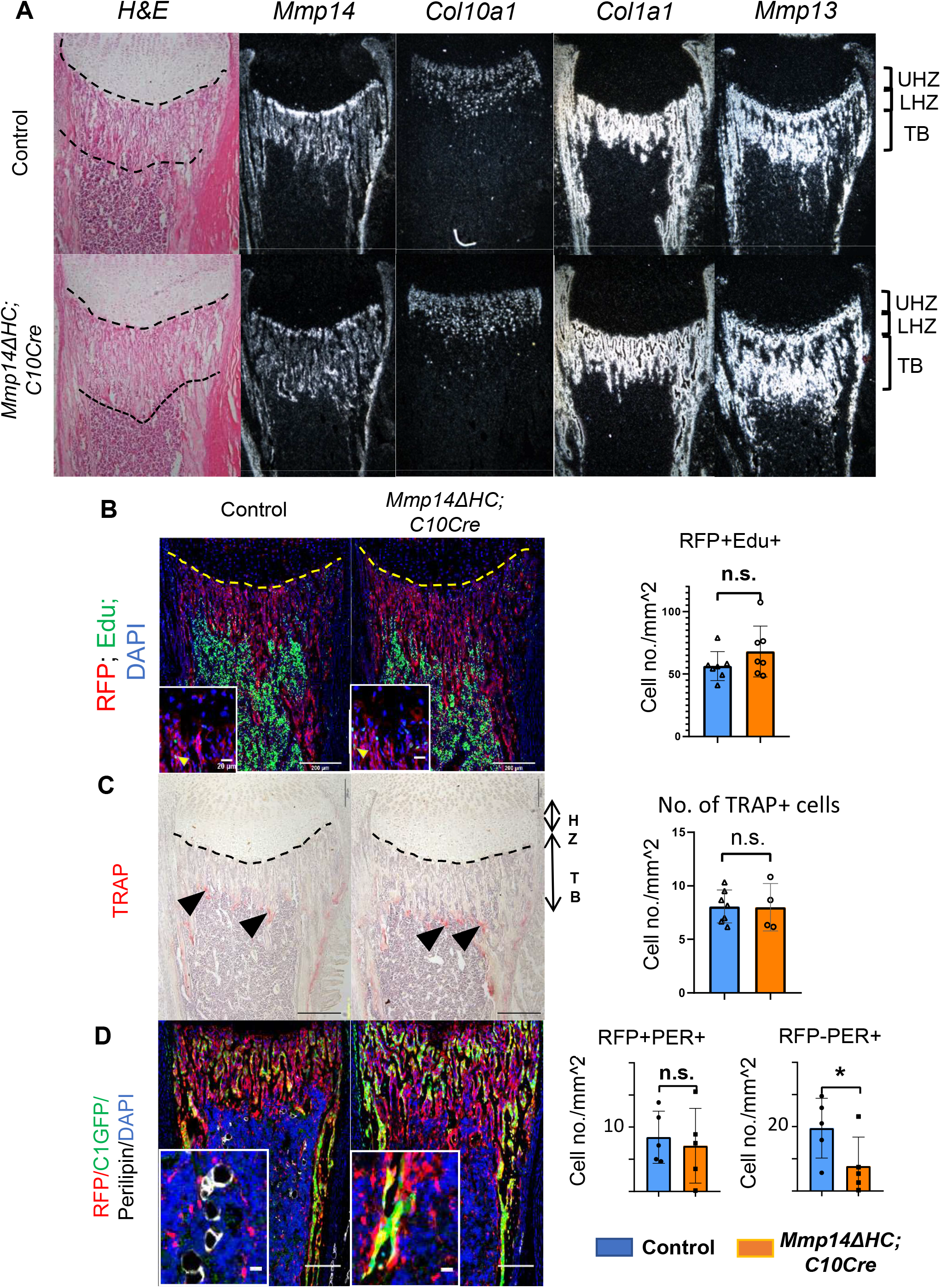
Molecular characterization of skeletal phenotype in *Mmp14^ΔHC^* mice. (A), H&E staining and in situ hybridization of *Col10a1*, *Col1a1*, *Mmp13*, *Mmp14* in *Mmp14^ΔHC^* and control mice at P10. White signals marks mRNA expression of corresponding genes. HZ, hypertrophic zone; TB, trabecular region. (B), Edu labeling assay in *Mmp14^ΔHC^* and control mice. HC-descendants, nucleus and proliferating cells were labeled with RFP(red), DAPI(blue) and Edu(green). (C), Tartrate-resistant acid phosphatase (TRAP) staining comparing *Mmp14^ΔHC^* and control mice. Black arrows mark TRAP+ cells HZ, Hypertrophic Zone. TB, Trabecular bone. Counting of TRAP+ cells suggests the number of osteoclasts are not significantly affected in the mutants(n=6 for control, n=4 for *Mmp14^ΔHC^* mutants). (D), Immunofluorescence staining of RFP, Perilipin, GFP and DAPI labeling tdTomato(Red), adipocytes(white), osteoblasts(green) and nucleus at P10. Quantitation of RFP+PER+, RFP-PER+, and PER+ cells at distal tibia comparing *Mmp14^ΔHC^* mutants and control at P10. Unpaired t-test are used for PER+ cells. Paired t-test is used for PER+RFP+ cells. Quantitation is shown in (n=5) (p<0.05) Unpaired t-test.

**Supplementary Figure 4.**
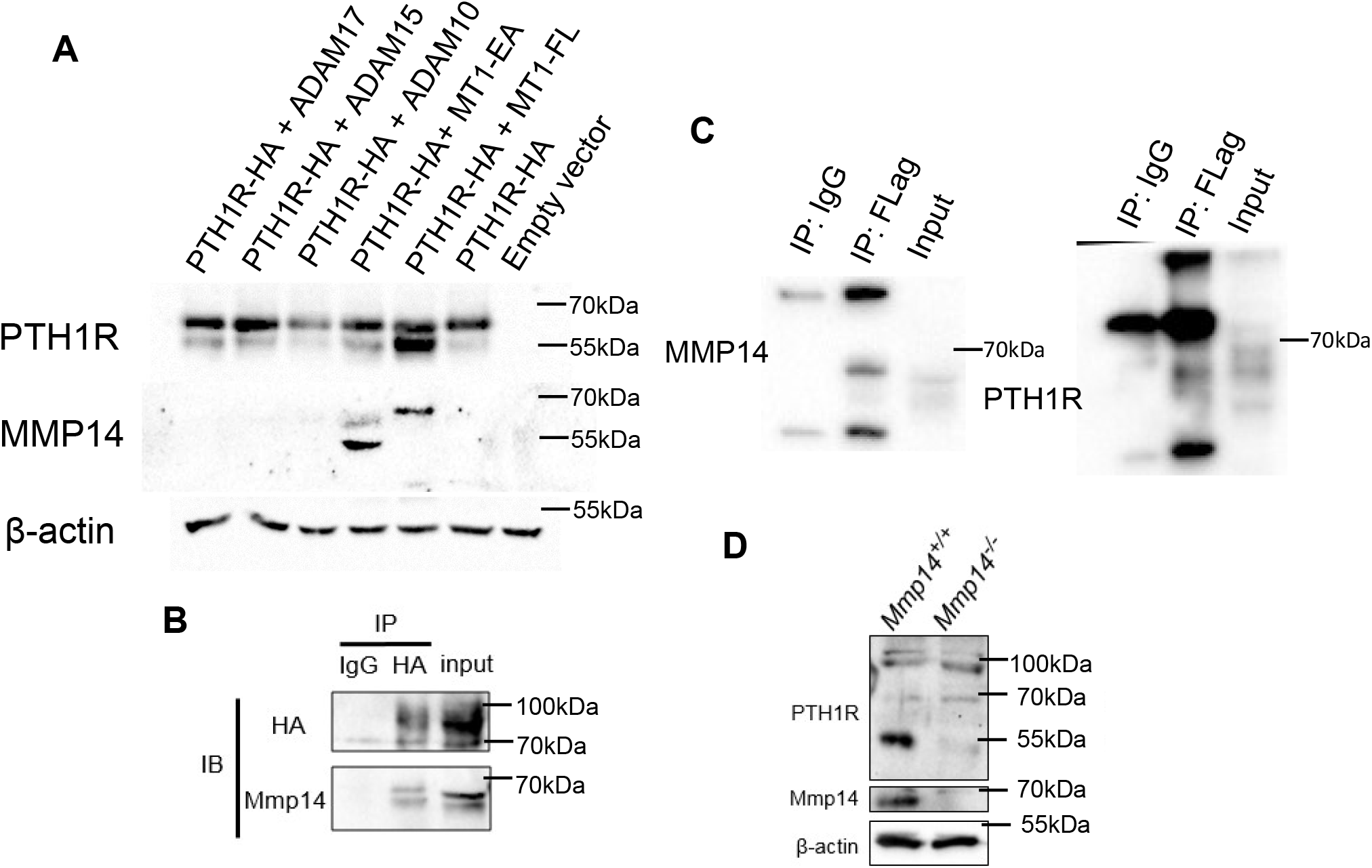
MMP14 is a major protease for PTH1R. (A), Western blot of HEK293T cells transfected with human PTH1R-HA, MT1-FL, MT1-EA, ADAM10, ADAM15 and ADAM17. (B), Co-immunoprecipitation of MMP14 with PTH1R in HEK293T cells transfected with MT1-FL and PTH1R-HA. PTH1R-HA was immunoprecipitated with anti-HA antibody in G-agarose. (C), Cell lysate from HEK293T cells transfected with PTH1R-HA, MMP14-Flag were immunoprecipitated with anti-Flag-tag antibody and analyzed by western blot. (D), Cleaved fragment of PTH1R were observed in cultured trabecular bone osteoblasts analyzed by western blot from Mmp14+/+(left lane) and Mmp14-/-(right lane) mice at P14.

**Supplementary Figure 5.**
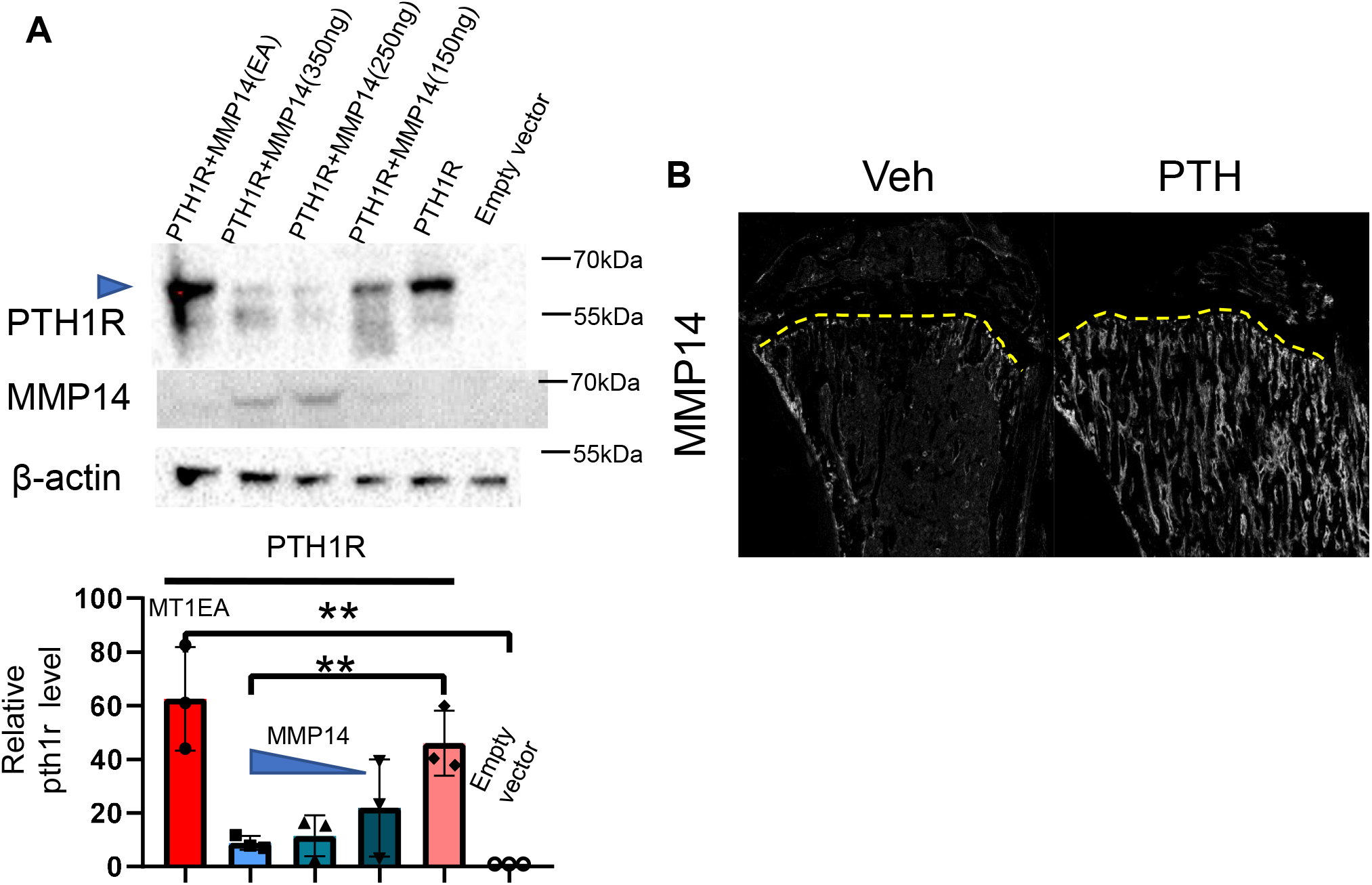
MMP14 destabilize PTH1R and can be upregulated by PTH. **(A),** Relative level of PTH1R in HEK293T cells co-transfected with empty vector, MMP14 and MMP14-EA. (**B**), Immunofluorescence staining of MMP14 in P56 mice treated with vehicle(left) and PTH(1–34).

**Supplementary Figure 6.**
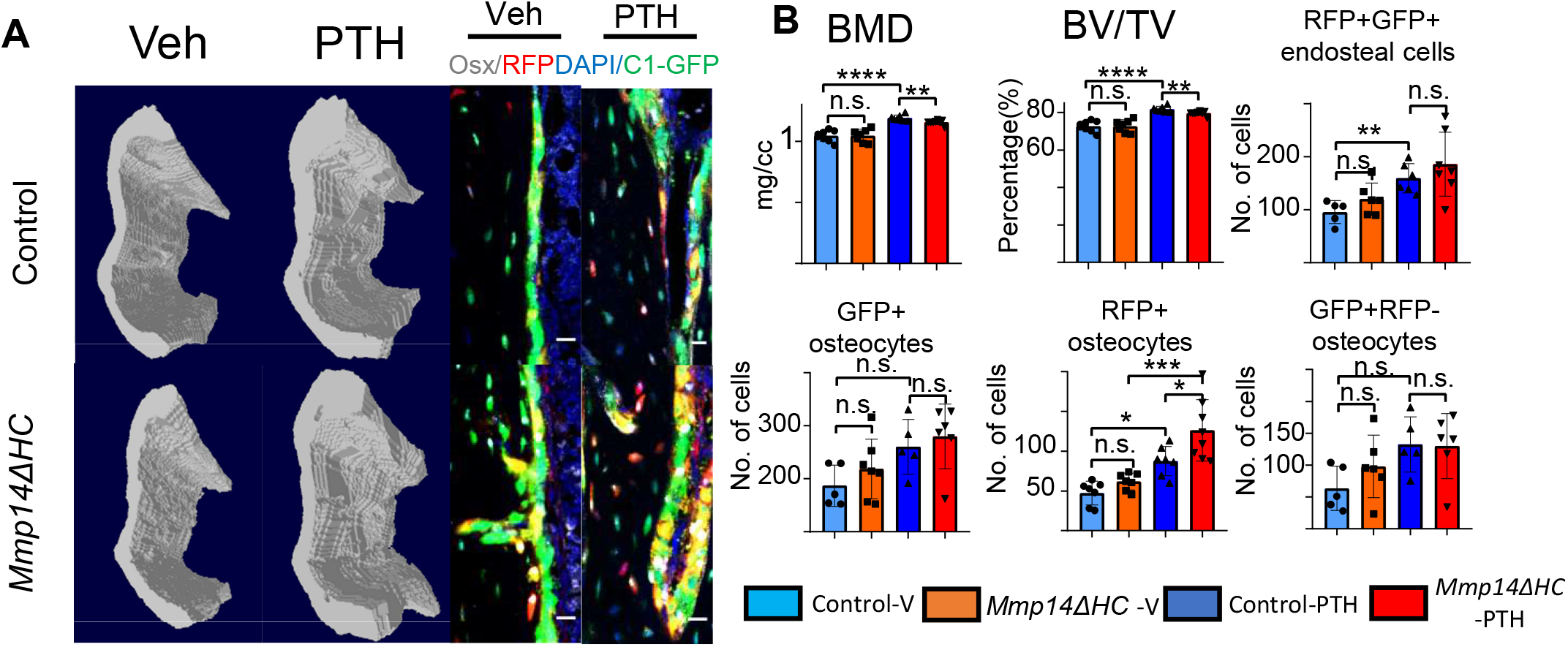
Differential response to PTH in cortical osteocytes compared to trabecular osteoblast. (**A**), Transverse images of cortical bone via MicroCT reconstruction in Control and *Mmp14ΔHC* mice treated with vehicle and PTH(1–34). Representative immunofluorescence images of OSTERIX, RFP, GFP and DAPI in cortical region are presented. (**B**), Quantitation of BMD, BV/TV in the cortical region as indicated in a. Quantitation of RFP+GFP+, RFP+, RFP-GFP+ osteocytes and endosteal RFP+ cells. * p<0.05, ** p<0.01, *** p<0.001 by two-tailed unpaired t-tests.

